# Single Nuclei Transcriptome Reveals Perturbed Brain Vascular Molecules in Alzheimer’s Disease

**DOI:** 10.1101/2021.12.28.474255

**Authors:** Özkan İş, Xue Wang, Tulsi A. Patel, Zachary S. Quicksall, Michael G. Heckman, Launia J. White, Laura J. Lewis-Tuffin, Kaancan Deniz, Frederick Q. Tutor-New, Troy P. Carnwath, Yuhao Min, Stephanie R. Oatman, Joseph S. Reddy, Minerva M. Carrasquillo, Thuy T. Nguyen, Charlotte C. G. Ho, Kimberly G. Malphrus, Kwangsik Nho, Andrew J. Saykin, Melissa E. Murray, Dennis W. Dickson, Mariet Allen, Nilüfer Ertekin-Taner

## Abstract

Blood-brain barrier (BBB) dysfunction is well-known in Alzheimer’s disease (AD), but the precise molecular changes contributing to its pathophysiology are unclear. To understand the transcriptional changes in brain vascular cells, we performed single nucleus RNA sequencing (snRNAseq) of temporal cortex tissue in 24 AD and control brains resulting in 79,751 nuclei, 4,604 of which formed three distinct vascular clusters characterized as activated pericytes, endothelia and resting pericytes. We identified differentially expressed genes (DEGs) and their enriched pathways in these clusters and detected the most transcriptional changes within activated pericytes. Using our data and a knowledge-based predictive algorithm, we discovered and prioritized molecular interactions between vascular and astrocyte clusters, the main cell types of the gliovascular unit (GVU) of the BBB. Vascular targets predicted to interact with astrocytic ligands have biological functions in signalling, angiogenesis, amyloid ß metabolism and cytoskeletal structure. Top astrocytic and vascular interacting molecules include both novel and known AD risk genes such as *APOE*, *APP* and *ECE1*. Our findings provide information on transcriptional changes in predicted vascular-astrocytic partners at the GVU, bringing insights to the molecular mechanisms of BBB breakdown in AD.

**Graphical Abstract:** Pericytes (yellow), endothelia (salmon) and astrocytes (purple) that form the gliovascular unit (GVU) at the blood brain barrier (BBB) were interrogated for their differentially expressed genes (DEG) and vascular cell (pericyte or endothelia) to astrocyte interactions using single nucleus RNA sequencing (RNAseq) transcriptome obtained from brains of Alzheimer’s disease (AD) patients and controls. We identified many upregulated (red) or downregulated (blue) DEGs in AD brains in these cell types. These genes have known biological functions in amyloid ß (Aß) clearance, immune modulation, astrogliosis and neuronal death. Novel predicted interactions were identified between vascular cells and astrocytic DEGs. Collectively, our findings highlight the vast transcriptome changes that occur at the GVU and provide mechanistic insights into BBB dysfunction in AD. This figure was created with Biorender.com.

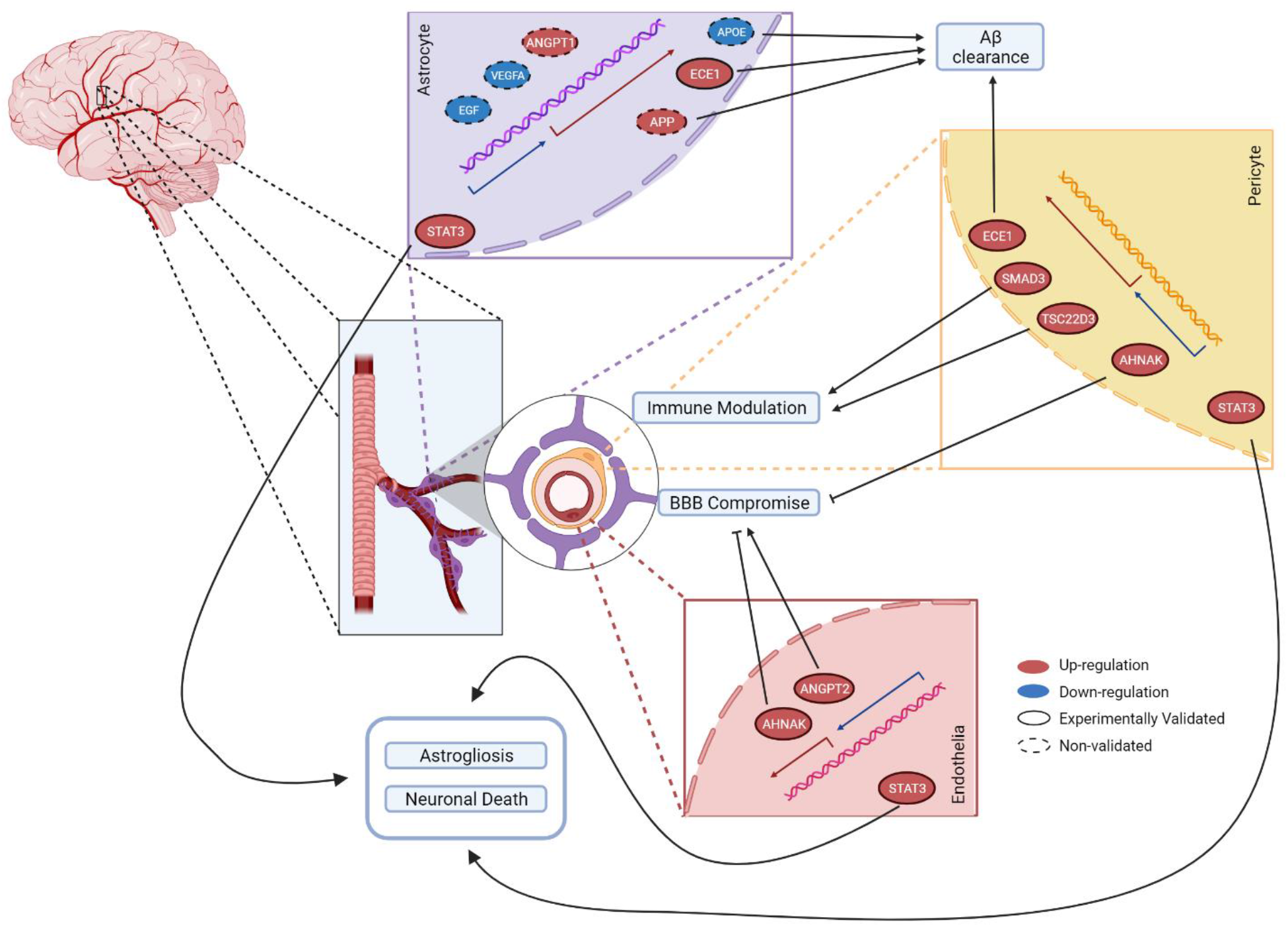

## Introduction

Single cell RNA sequencing (RNAseq), a technology which enables researchers to obtain transcriptome of individual intact cells (scRNAseq) or nuclei (snRNAseq)^1,2^, has been utilized to identify different cell types and/or subtypes in Alzheimer’s disease (AD) and healthy brains, to identify cellular states and cell activation, to describe vulnerable cell populations, and to elucidate perturbed genes and pathways in each cell type in AD^3–10^.

To date most single cell transcriptome studies of AD brains focused on neuronal cells and more abundant glial cells, with relatively little known on transcriptional changes in vascular cells, namely endothelia and pericytes, and their interaction with other central nervous system (CNS) cells, despite their important roles in AD progression^11,12^. Mathys et al. reported differentially expressed genes for neurons, astrocytes, oligodendrocytes, oligodendrocyte progenitor cells (OPCs), and microglia, highlighting myelination pathways as being significantly disturbed in dorsolateral prefrontal cortex (DLPFC)^13,14^, as previously shown in bulk RNA sequencing (RNAseq) studies^13,15^. Zhou et al. surveyed transcriptional changes of microglia, oligodendrocytes and astrocytes in DLPFC of human AD brain and a mouse model^8^. Kun et al. reported the selective vulnerability of neuronal subpopulations and reactive astrocytes in caudal entorhinal cortex and the superior frontal gyrus of AD brains^10^. To our knowledge the only single cell level study of vascular cells in AD patient brains is by Lau et al., who reported endothelial cell transcriptome changes in AD which were related to angiogenic functions^9^.

The gliovascular unit (GVU) is the multicellular functional unit of the central nervous system (CNS) that consists of endothelia, pericytes, and astrocytes, which comprise the blood brain barrier (BBB)^16–18^. Impairment in BBB is a key feature in AD, which leads to entry of neurotoxic blood derived products, cells and pathogens associated with inflammatory response and reduced cerebral blood flow^11^. Accumulation of amyloid ß (Aß) deposits around cerebral vasculature is thought to be both a cause and consequence of this BBB impairment^19,20^. Despite the evidence that BBB impairment contributes to AD pathogenesis and progression^21,22^, precise transcriptional changes in the GVU in AD, molecular interactions between brain vascular cells and astrocytes, and the correlations of their transcriptome changes with AD neuropathology at the single cell level remain to be established.

In this snRNAseq study of temporal cortex samples from 12 AD and 12 unaffected elderly control brains (**Figure 1, Supplementary Figure S1, Supplementary Table S1**), we focused on the transcriptional landscape of the GVU and its changes in AD. We report the discovery of three distinct vascular and three astrocytic clusters, characterize their molecular signatures, enriched biological processes and transcriptome changes in AD. We further identify predicted brain vascular-astrocyte molecular interaction partners for each vascular and astrocytic cluster using a knowledge-based algorithm^23^. We prioritize the vascular genes for their strength of astrocytic connections and differential expression in AD, validate their brain expression changes and demonstrate their associations with AD neuropathologies, as well as age and sex. Our findings demonstrate that most differentially expressed brain vascular genes in AD are upregulated and reside in the activated pericyte nuclei cluster. The upregulated activated pericyte cluster genes are enriched for hormone receptor binding processes and include signalling molecules such as *SMAD3* and *STAT3*, which have strong astrocytic ligands interactions. Top astrocytic and vascular interacting molecules with differential expression in AD brains include known AD risk genes such as *APOE*, *APP* and *ECE1*, as well as novel genes. Our findings provide information on brain vascular expression changes at a single nucleus level in AD and uncover new vascular-astrocytic interactions that may lead to breakdown of the GVU and influence propagation of AD neuropathology.

**Figure 1:**
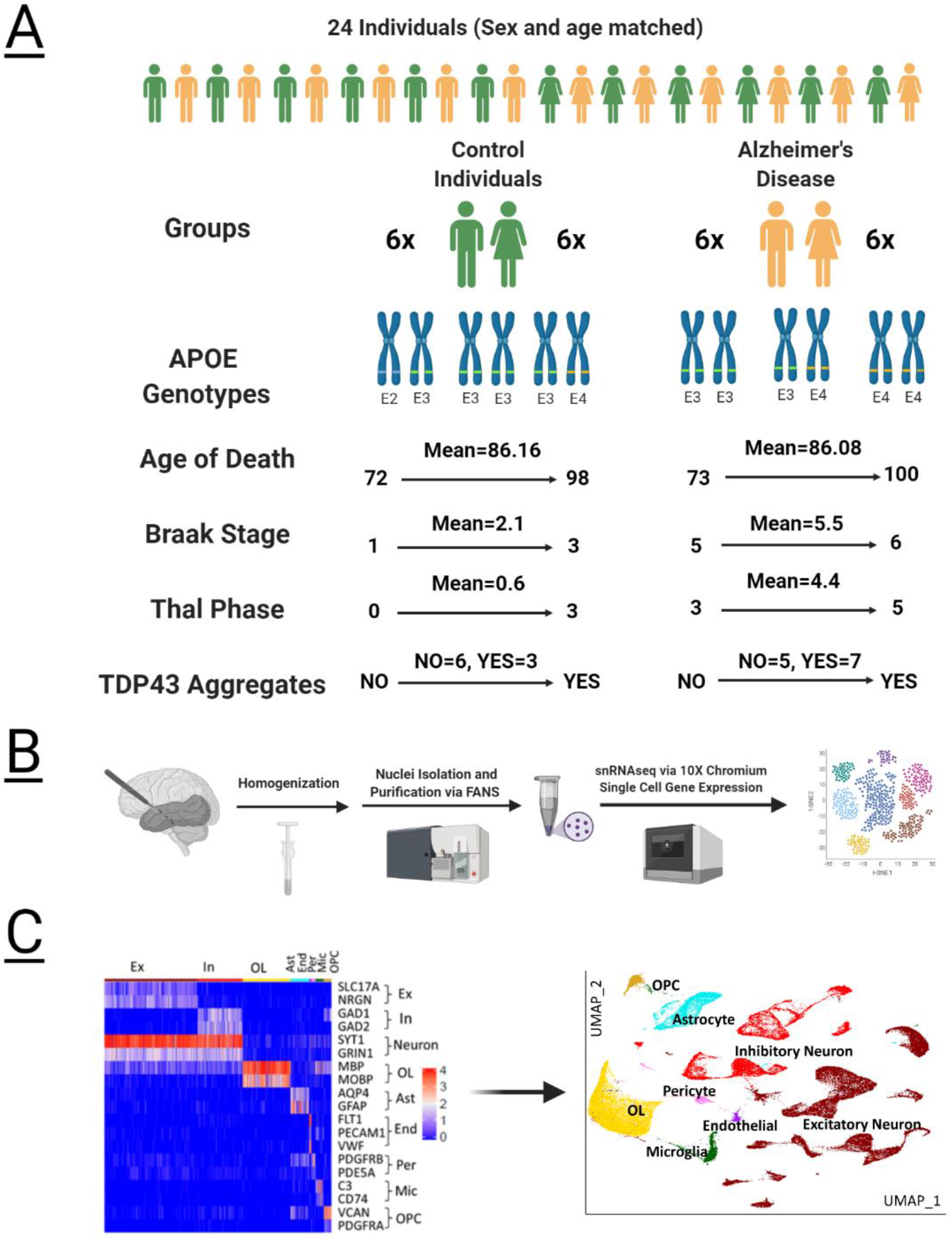
Summary of the snRNAseq approach utilized in this study. A) Post-mortem temporal cortex tissue from 24 Individuals that comprise sex and age matched AD and control individuals were used in this study. B) Development and optimization of nuclei isolation protocol for snRNAseq platform. C) Well-established cell type markers were used to annotate nuclei clusters. This figure was created with Biorender.com.

## Results

### Development of an unbiased snRNAseq approach to survey brain transcriptome of all cell types

Purity and quality of isolated nuclei from human brains are critical for optimal snRNAseq results. Most nuclei isolation methods rely on sucrose gradient^24^ or fluorescence-activated nuclear sorting (FANS)^25^ to obtain high quality and pure nuclear fraction; but these approaches can limit detection of rare cell types. We developed a nuclei isolation method that allows detection of all cell types with high purity with limited bias against rarer cell types (**Supplementary Figures S2-S3**). We selected 12 neuropathologic AD and 12 age- and sex-matched control patients (**Figure 1a, Supplementary Figure S1, Supplementary Table S1**). We obtained snRNAseq profiles from their temporal cortex (TCX) using 10x Genomics platform, which yielded 87,493 single nucleus gene expression profiles (**Figure 1b, Supplementary Table S1**).

We applied quality control (QC) steps based on number of genes and unique molecular identifiers (UMIs) per nuclei (**Supplementary Figure S4**), resulting in 79,751 high quality nuclei in 35 clusters that were annotated for their types according to published cellular markers^26^. Heatmap visualization using well-established cell type markers further confirmed the cell type assignment for the clusters (**Figure 1C**). All clusters include nuclei from > 20 individuals, i.e. > 80% of cohort, except the two smallest clusters which contain 161 and 159 nuclei (**Supplementary Tables S2-S3**). Clusters represent eight cell types (**Figure 1C**) as follows: 14 excitatory neuronal (41% nuclei), 9 inhibitory neuronal (20%), 3 oligodendrocytic (21%), 3 astrocytic (8%), 1 endothelial (1%), 2 pericyte (2%), 2 microglial (3%) and 1 oligodendrocyte progenitor nuclei clusters (3%).

We analyzed each cluster to determine whether diagnosis, age, sex, and pathology scores have an impact on their nuclei proportions (**Supplementary Figures S5-S6, Supplementary Tables S4-S6**). For most clusters, there was no statistically significant difference in nuclei proportions by these variables with the exceptions of excitatory neuronal cluster 16 (cl.16) that has a lower proportion of cells in AD and also negatively associated with Braak and Thal, as expected; inhibitory neuronal cl.10 also negatively associated with Braak; astrocytic cl.9 positively associated with presence of TDP-43 and astrocytic cl.26 with lower proportion in females.

### Unique AD associated differentially expressed genes (DEGs) and pathways are detected in distinct brain vascular clusters

Three vascular nuclei clusters were identified – cl.25, cl.27 and cl.31, containing 943 (AD:430, control:513), 765 (AD:326, control:439), and 502 (AD:245, control: 257) nuclei respectively (**Figure 2A, Supplementary Figure S7**). All three clusters express BBB-specific transcription factor *LEF1^27^* (**Figure 2B, Supplementary Figure S7B**), which is not expressed in any other clusters. Smooth muscle cell markers of peripheral vasculature *ACTA2*, *RBP1^28^*, *SMN1^29^* have essentially no expression in any of the brain vascular clusters (**Supplementary Figure S7**).

**Figure 2:**
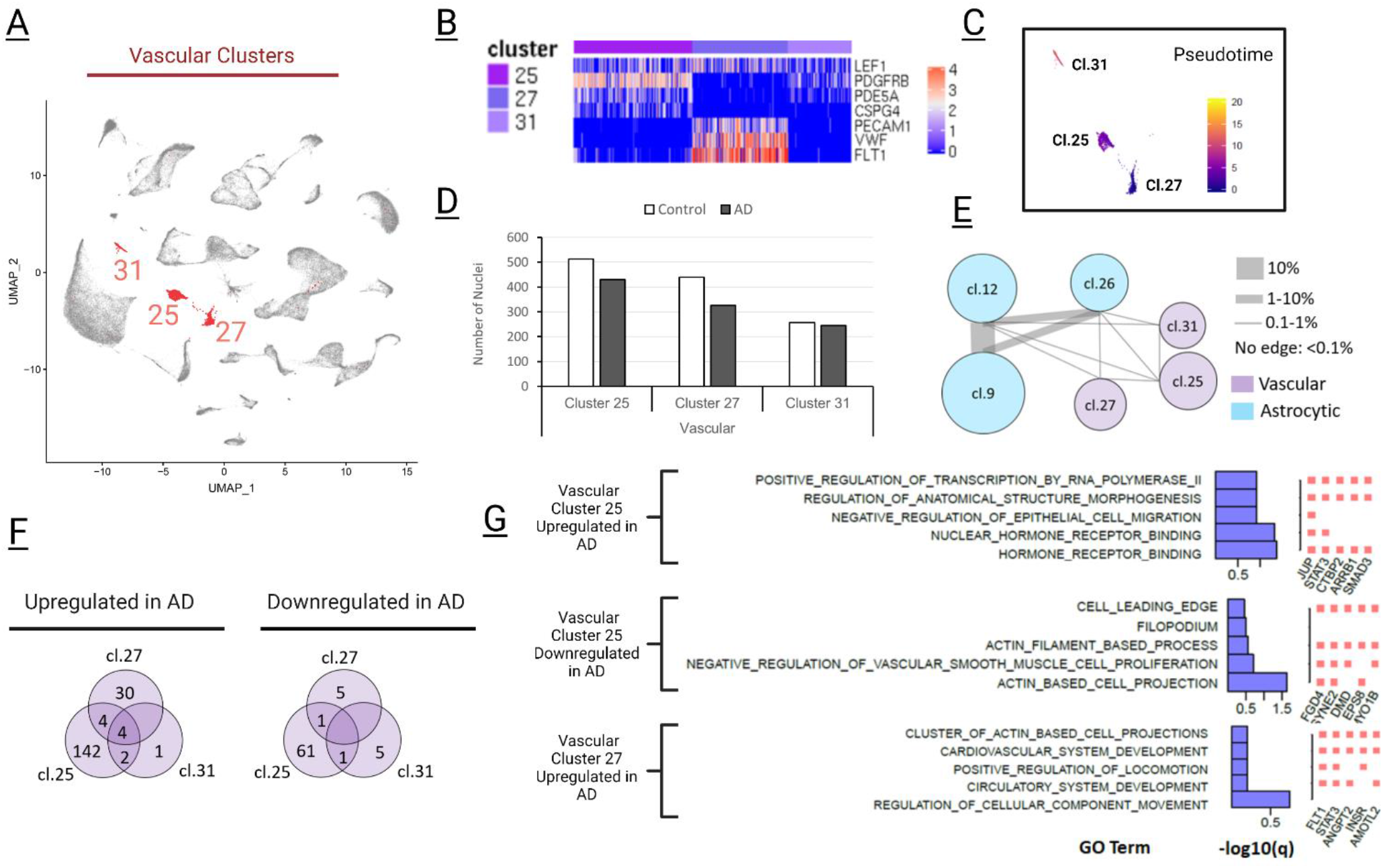
Nuclei in vascular clusters demonstrate altered genes and pathways in AD. A) Three vascular clusters were demonstrated in UMAP plots. B) While cluster 27 demonstrated expression of endothelial signature genes, Clusters 25 and 31 showed upregulation in pericyte marker genes. C) Pseudotime analysis of the three vascular clusters displays their different expression profiles. D) Proportions of AD and control nuclei in these clusters were not significant. E) The constellation plot displays the relatedness of the 3 vascular and 3 astrocyte (**see Figure 3**) clusters, based on post-hoc classification of cells. The thickness of the connecting line between any two clusters was determined by the percent of cells that are ambiguously assigned. Astrocytes clusters demonstrated greater relatedness as shown by the thick connecting lines (~1-10%). Vascular clusters, on the other hand, demonstrated more distinct cell populations with thin connecting lines (~0-1%). F) Differentially expressed gene (DEG) profile in AD were different between these clusters as very few DEGs were shared among these clusters. G) Top GO Term Enrichment analysis were summarized for activated pericyte cluster cl.25 and endothelia cluster cl.27. GO Enrichment Term analysis displayed several upregulated and downregulated pathways in AD for cl.25 and upregulated pathways in cl.27.

The three vascular clusters are well separated in the reduced dimension UMAP plot (**Figure 2A**) and in pseudo-time plot (**Figure 2C**). Furthermore, using random forest classification to identify any ambiguous or intermediate cells between each of the two clusters, we determined that only <0.5% of the cells were ambiguous (**Figure 2E, Supplementary Table S7**), further highlighting their distinct gene expression. None of the vascular clusters demonstrated any bias in their distributions across either diagnostic group, pathology scores, sex, age or *APOE* (**Figure 2D, Supplementary Figure S6, Supplementary Tables S4-6**).

To understand their unique biological functions, we identified signature genes that were detected in at least 50% cells in the cluster of interest, had average expression >= 2.0X that of either of the other two clusters with Bonferroni-corrected p-value < 0.05. This resulted in 89, 147 and 64 signature genes for cl.25, cl.27 and cl.31 respectively (**Supplementary Table S8**), using which we performed GO enrichment analyses (**Supplementary Figure S8**, **Supplementary Tables S9-11**).

Cl.27 is enriched in endothelial GO terms (**Supplementary Figure S8, Supplementary Table S10**) and highly expresses endothelial markers *VWF*, *FLT1* and *PECAM1/CD31^30,31^*,endothelial damage associated genes such as *VWF^32^*, *ABCG2^33^*, *ABCB1^34^* and angiogenesis associated genes such as *ENG^35^*, *TGM2^36^*, and *ERG^37^* in comparison to the other two vascular clusters. In contrast, cl.25 and cl.31 have expression signatures consistent with pericytes. Compared to endothelial cl.27, these two clusters have upregulation of pericyte markers *PDGFRB^38,39^* and *PDB5A^30,40^* (**Figure 2B**). Additionally, cl.25, but not cl.31, displays activated pericyte phenotype as it has upregulation of *CSPG4/NG2^39,41^*, *RGS5^42^*, *and P2RY14^43^*. Further, cl.25 has high expression of genes related to nutrient and ion transport (*SLC12A7^44^*, *SLC6A12^45^*, *SLC19A1^46^*), and formation of blood brain barrier (*COL4A1^47^*, *CDH6^48^*, *SNTB1^49^*) in comparison to cl.27 and cl.31. Finally, cl.31 expression profile is consistent with pericytes in basal membrane that reshape extracellular matrix (ECM) by relative increased expression of collagens (*COL1A2*, *COL12A1*, *COL15A1*), fibrillin (*FBN1*) and laminins (*LAMA2*, *LAMB1*, *LAMC1*). Indeed, the top cl.31 signature gene GO terms are related with the ECM (**Supplementary Figure S8, Supplementary Table S11**). This cluster also displayed elevated expression of angiogenesis related genes such as *FLVCR2^50^*, *ANTXR2^51^* and *PRRX1^52^*, when compared with cl.25 and cl.27. In summary, based on their unique expression profiles, cl.27, cl.25 and cl.31 are named endothelial, activated pericyte, and resting pericyte clusters, respectively.

We next performed DEG analyses for each vascular cluster to compare their gene expression in AD vs. control tissue using MAST R package^53^. Imposing q value < 0.05, absolute log (fold change) > 0.1 and detection of gene expression in >= 20% cells, 215 (152 up, 63 down), 44 (38 up, 6 down), and 13 (7 up and 6 down) DEGs were identified in cl.25, cl.27 and cl.31 respectively (**Figure 2F, Supplementary Table S12**). Four genes are up-regulated in AD across all three vascular clusters (*INO80D*, *LINGO1*, *RASGEF1B*, *SLC26A3*) and no down-regulated genes are shared. Most DEGs were detected in cl.25, supporting this cluster being activated in AD. The limited number of overlapping DEGs in any two clusters (**Figure 2F**) further confirmed that these clusters are distinct from each other and likely have different biological roles. GO enrichment analyses were performed for those vascular cluster DEGs that had sufficient number of DEGs, i.e. genes up or down in AD in cl.25, and genes up in cl.27 (**Supplementary Tables S13-15**). The top 5 GO terms and their frequently observed DEGs are shown in **Figure 2G**. These top genes are growth factor related and upregulated in activated pericyte cl.25 or endothelial cl.27 clusters (*FLT1*, *SMAD3*, *STAT3*, *INSR*), angiogenesis related and upregulated in endothelial cl.27 (*ANGPT2*, *AMOTL2*) and cytoskeleton related and downregulated in activated pericyte cl.25 (*DMD*, *FGD4*, *MYO1B*). These findings support AD-related expression changes in distinct brain vascular cells.

### Astrocyte clusters have AD associated differentially expressed genes (DEGs)

Given the known role of astrocytes in the GVU of the BBB^16–18^, we next focused on the astrocytic clusters and their DEGs associated with AD. Three astrocytic clusters cl.9, cl.12 and cl.26 were identified, encompassing 3129 (AD:1716, normal:1413), 2597 (AD:1310, normal:1287) and 815 (AD:462, normal:353) cells, respectively (**Figure 3A**). Cl.12 has features of proliferative reactive astrocytes with high expression of *GFAP*, *CD44*, *VCAN* and low expression of *NDRG2^54^* (**Figure 3B**). Unlike the vascular clusters, the astrocytic clusters were less distinct as illustrated by their pseudotime analysis **(Figure 3C)** and by their percentage of ambiguous or intermediate cells (1-10%, **Supplementary Table S7, Figure 2E**). Signature genes were identified for astrocytic clusters as done for the vascular clusters. There were 4, 8 and 124 signature genes for cl.9, cl.12 and cl.26 respectively. GO term analysis could only be conducted for cl.26 signature genes which showed enrichment for ion transportation, synaptic signaling and myelination terms (**Supplementary Table S17**), suggesting that this may either be a mixed cluster or one involved in astrocyte-neuron and oligodendrocyte interactions.

**Figure 3:**
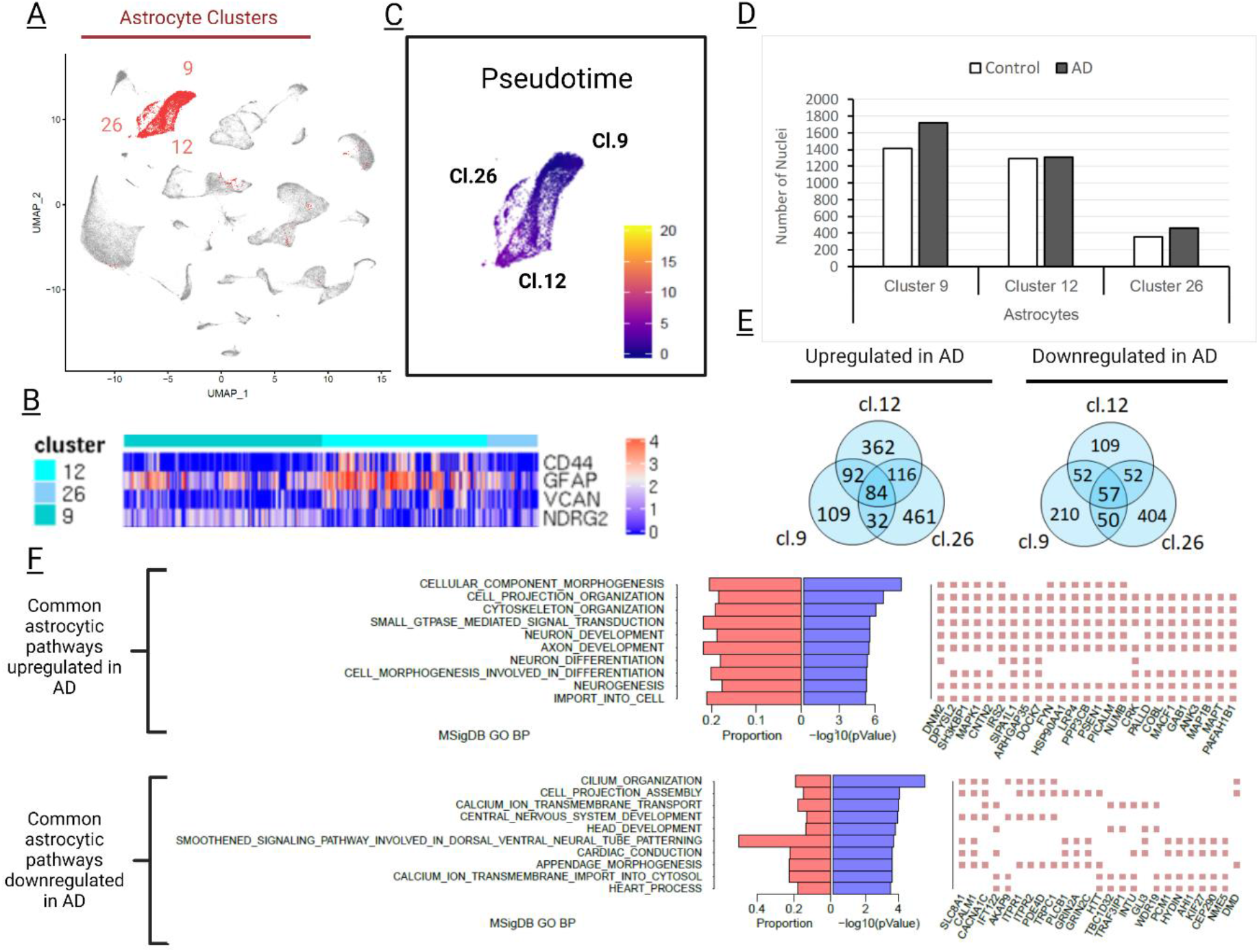
Transcriptome changes in AD astrocytes. A) Three different astrocyte clusters were observed as shown in the UMAP plot. B) Despite similarities of these clusters, astrocytic cluster 12 displays a more reactive profile because of high expression of *CD44*, *GFAP*, *VCAN*, and low expression of *NDRG2*. C) Pseudotime analysis demonstrated that these clusters are similar in their transcriptomics profile. D) These clusters did not show significant difference in the nuclei proportions according to diagnosis group. E) Despite similarities in their transcriptome, these clusters also had unique DEGs. F) GO Term enrichment analysis revealed biological pathways harboring DEGs commonly affected by AD across all astrocytic clusters.

As for the vascular clusters, the proportion of AD and control nuclei in astrocytic clusters were not statistically different (**Figure 3D, Supplementary Figure S6, Supplementary Table S4**). However, unlike vascular clusters, astrocyte cluster 26 demonstrated significantly higher number of nuclei from male brains **(Supplementary Table S5)**. The only other nuclei proportion difference was for cl.9, which showed a higher proportion of TDP43 positive cells (**Supplementary Table S6**).

Genes in astrocytic clusters that were differentially expressed between AD and control brains were identified. Cl.9, cl.12 and cl.26 contain 686 (317 up, 369 down), 924 (654 up, 270 down), 1256 (693 up, 563 down) DEGs, respectively (**Figure 3E, Supplementary Table S18**). Top 5 GO terms of DEGs upregulated in AD within cl.9 and cl.12 include actin cytoskeleton, cell differentiation related terms, whereas for cl.26 the top enriched term in upregulated DEGs are cytoskeleton, neurogenesis, and ensheathment of neurons (**Supplementary Tables 19-21**). For DEGs that are downregulated in AD in the astrocytic clusters, the top enriched GO terms include cell signaling, neurogenesis and cilia/motility related processes (**Supplementary Tables 22-24**). Unlike vascular DEGs, about 18% of the astrocytic DEGs are shared in two or more clusters. GO enrichment analyses of these genes commonly perturbed in AD astrocytes demonstrate enrichment of cytoskeleton and neurogenesis related terms for upregulated; and cilium and calcium transport related terms for downregulated genes (**Figure 3F**). Comparison of the astrocytic cluster DEGs in our study to a previously published study that focused on astrocytes^9^ revealed significant overlap, as well as unique genes (**Supplementary Figure 9**). Our findings support widespread transcriptome perturbations in AD astrocytes which have many shared DEGs between their clusters that have more similar transcriptional profiles for this cell type in comparison to the brain vascular clusters.

### Ligand-target interactions between astrocytic and vascular AD-associated genes

Cell biology studies of the GVU have discovered multiple interactions between astrocytes and brain vasculature that are mediated through ligand-target interactions, although systematic efforts are needed to discover the vast and complex molecular relationships between these cells of the BBB^16–18^. To systematically identify a prioritized set of vascular targets that are regulated by astrocyte ligands and consequently influence brain vascular functions at the GVU of the BBB, we used transcriptome data from the brain vascular and astrocyte clusters and the Nichenet^23^ analytic platform that utilizes prior knowledge of such interactions. As our goal was to determine those vascular target-astrocyte ligand pairs that are most perturbed in AD, we confined our analyses to the significant vascular and astrocytic DEGs. Using significant astrocytic DEGs and Nichenet^23^, we identified 22 potential ligands in cl.9, 23 in cl.12 and 33 in cl.26, respectively (**Supplementary Table S25**). Some of these potential ligands were identified in more than one astrocytic cluster, thus resulting in a combined pool of 59 ligand genes (**Supplementary Figure S10**). To prioritize for corresponding targets in each vascular cluster that had the strongest analytic evidence, we selected those that had a ligand in each astrocytic cluster. Using this criterion and taking all significant DEGs of activated pericyte cl.25, endothelial cluster cl.27 and resting pericyte cluster cl.31, we identified 19, 6 and 1 potential vascular targets, respectively, corresponding to one or more astrocytic ligands, resulting in 24 target genes (**Supplementary Figure S11, Supplementary Table S25**).

These 24 brain vascular target candidates include genes with diverse biological functions (**Figure 4A**), including cytoskeleton and ECM-related (*TIMP3^55^*, *DMD^56^*, *AHNAK^57^*, *SLC38A2^58^*, *STARD13^59^*); growth factor-related (*STAT3*, *SMAD3*, *TGFB1*, *TFPI*, *EGFR*)^60–62^; glucocorticoid-related and anti-inflammatory (*NR3C1*, *TSC22D3*)^63^; angiogenesis (*AMOTL2*, *ANGPT2*)^64,65^; as well as *ECE1*, an AD-related gene, that is involved in Aβ clearance and vasoconstriction^66,67^. There were established AD genes amongst the top astrocytic ligands namely *APOE* corresponding to predicted vascular target *TSC22D3* with high estimated regulation strength and *APP* with high regulation strength for *ECE1*. (**Figure 4A, Supplementary Table S25**).

**Figure 4:**
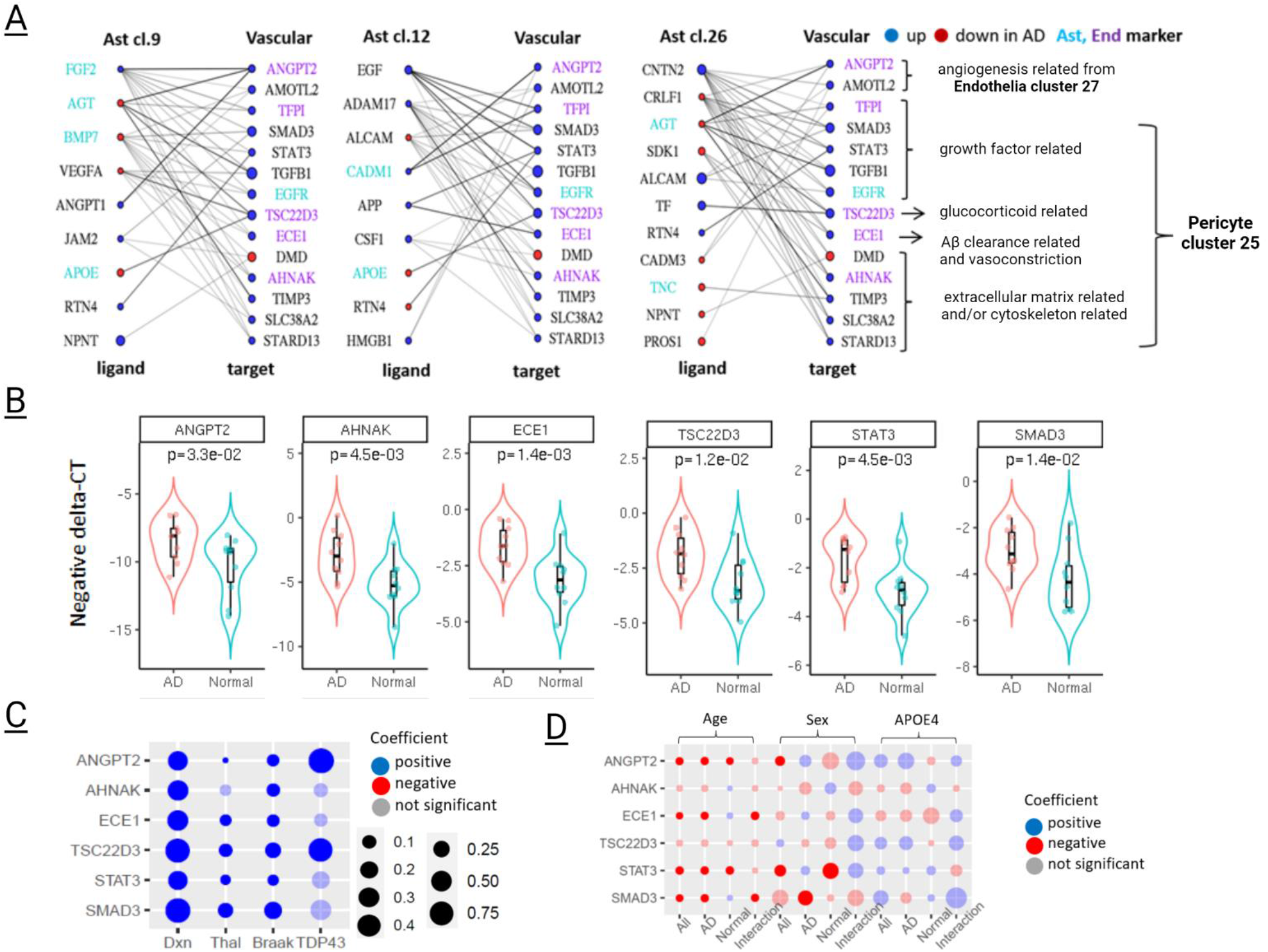
Interaction of predicted brain vascular targets and their astrocyte ligands. A) Strength and direction of NIchenet vascular target-astrocyte ligand interactions. Left side columns: predicted ligands in astrocyte clusters. Right side columns: predicted targets in vascular clusters. Edge: potential regulation between ligands and target genes, the thickness of which reflects the strength of regulation according to NicheNet prior model. Cell type marker genes in literature were colored as cyan for astrocyte and purple for endothelial genes. Direction of change in AD is denoted as blue for up and red for downregulation. Known biological functional categories for the vascular targets are shown to the right of the figure. B) Expression of six prioritized vascular target genes were validated via qPCR from nuclear fractions. Distribution of negative delta-CT from qPCR experiment of six prioritized vascular target genes, which reflect the expression of these gene. One-sided Wilcoxon rank sum tests were performed to test whether these genes were expressed higher in AD compared to control nuclei, as observed from snRNAseq. C) The association between AD-related pathologies and snRNAseq expression of six vascular genes. Blue bubble: significant positive association with pathologies at q-value≤0.05. Red bubble: significant negative association with pathologies at q-value≤0.05. Gray-masked bubble: not significant. Size of the bubble refects the association coefficient. D) The association between age at death, sex and *APOEε4* and expression of six prioritized vascular genes in combined AD and controls, ADs only and controls only, respectively, as shown in the first three columns for each association. The fourth column for each association shows the interaction effects between age, sex or *APOEε4* with diagnosis on gene expression. A significant interaction effect indicates that the association between the risk factor and gene expression is dependent on disease status. Blue bubble: significant positive association at q-value≤0.05. Red bubble: significant negative association at q-value≤0.05. As males or *APOEε4* carriers were denoted as 1 whereas females or *APOEε4* non-carriers as 0, blue means higher in males or *APOEε4* carriers and red means lower in males or *APOEε4* carriers. Gray-masked bubble: not significant. Size of the bubble refects the association coefficient.

We next prioritized the 24 predicted vascular targets with regulatory astrocytic ligands, for validations of their expression levels by quantitative PCR (qPCR). We selected genes representative of the biological functions determined amongst these 24 genes (**Figure 4A, Supplementary Table S26**). For validations, we chose *SMAD3* and *STAT3* that are growth-factor related signalling molecules*^61,68,69^*. *SMAD3* has the strongest predicted interactions with astrocytic ligands amongst all 24 vascular genes, is upregulated in activated pericyte cluster cl.25 (**Supplementary Table S26**) and is one of the most frequently observed genes in the GO terms enriched for this cluster (**Figure 2G**). *STAT3*, also a strong vascular target, is upregulated in both cl.25 and endothelial cl.27 (**Figure 2G, Supplementary Table S26).** Other selected genes were *AHNAK*, *ANGPT2*, *ECE1* and *TSC22D3. AHNAK*, the second most strongly connected vascular target encoding a structural protein involved in BBB integrity^57^, is an upregulated DEG in activated pericyte cl.25 **(Figure 4A, Supplementary Table S26)**. *TSC22D3* and *ECE1* that are also upregulated in activated pericytes have known Aß-related functions^66,67,70^, predicted astrocytic ligands that are AD genes (**Figure 4A**), and anti-inflammatory and vasoconstrictive properties, respectively. Finally, *ANGPT2* that is the most significantly upregulated DEG in endothelial cl.27, is a strong target in this cluster (**Figures 2G, 4A, Supplementary Table S26**). *ANGPT2* is involved in angiogenesis^65^ and like *SMAD3* and *STAT3*, is a signalling molecule.

Using nuclei isolated from the same brain region (temporal cortex) of the same cohort, we measured the expression of these genes using qPCR, which validated their significant differential expression in AD vs. control brains as observed in our snRNAseq data (**Figure 4B, Supplementary Table S27**). For these six experimentally validated genes, we further tested their expression associations with Thal score for amyloid-beta pathology, Braak stage for tau pathology and TDP-43 pathology (**Figure 4C**). For *ANGPT2*, expression in endothelial cl.27 is used, whereas for other genes expression in activated pericyte cl.25 is used, per their Nichenet results. As expected, all 6 vascular target genes that are higher in AD brains also have higher expression with increasing Braak stages and Thal phase (**Figure 4C**). These associations are statistically significant (q<0.05) with the exception of Thal phase association with AHNAK levels (q=0.13). All genes are also higher in the presence of TDP-43 pathology with *ANGPT2* and *TSC22D3* reaching significance (q<0.05).

There are also significant associations with age and sex, but not *APOEε4*. Brain levels of *ANGPT2*, *ECE1*, *STAT3* and *SMAD3* are reduced with aging (**Figure 4D**). These associations are driven by the AD group for *ECE1* and *SMAD3*, which have significant diagnosis by age interaction for expression associations, but not the other genes. Regarding sex differences, *ANGPT2* and *STAT3* expression is significantly lower in males in the combined group, which seems to be driven by controls especially for latter. *SMAD3* is lower in AD males. Collectively, these results demonstrate that while all 6 vascular targets are higher in AD temporal cortex and with AD-related pathologies, the levels of some of these genes are also influenced by age and sex in a differential manner by diagnosis.

## Discussion

Single cell and single nuclei approaches have been instrumental in revealing the molecular perturbations in AD, however most of these studies have been focused on relatively common brain cell types^4–6,8^. In our study, we have analyzed temporal cortex brain tissue from 12 AD and 12 control donors using snRNAseq and obtained 79,751 post-QC nuclei, of which 4,604 were classified as brain vascular cells. Despite the known breakdown of BBB in AD^11,19,22,71,72^, there is relative paucity of sn/scRNAseq studies focusing on brain vascular cells in AD, likely due to their low frequency, with the exception of Lau et al.^9^ study which obtained gene expression profiles from ~2,400 endothelial nuclei in 12 AD and 9 control brain samples.

Our snRNAseq data from brain vascular nuclei enabled the following observations. First, we identified three distinct vascular clusters which could be classified as activated pericytes (cl.25), endothelia (cl.27) and resting pericytes (cl.31), owing to the unique expression profiles of their highly expressed signature genes. Second, we identified DEGs in these vascular clusters the largest numbers of which were in the activated pericyte cl.25 (152 up, 63 down), followed by endothelial cl.27 (38 up, 6 down) and resting pericyte cl.31 (7 up, 6 down). The limited number of overlapping DEGs amongst the brain vascular clusters underscore their distinct nature. Third, enriched GO terms amongst the vascular cluster DEGs highlight perturbed biological processes in brain vasculature where upregulated activated pericyte cl.25 had many signalling molecules such as *SMAD3^61^* and *STAT3^61,68,69^*, whereas downregulated genes in this cluster had cytoskeletal genes such as *DMD^56^*, with enrichment for hormone receptor binding and actin-based processes, respectively. Endothelial genes (cl.27) upregulated in AD include angiogenesis related genes such as *ANGPT2^65^* and *AMOTL2^64^*. These findings demonstrate the vast transcriptional changes in brain vascular cells in AD and nominate molecules and pathways that may propagate the known BBB dysfunction and breakdown in this condition^73,74^.

Given the known interactions and proximity between astrocytes and brain vascular cells, i.e. endothelia and pericytes, at the BBB^11,75^, we sought to discover those molecules that have strong interactions between these key cell types of the GVU. For this purpose, we used the analytic approach of Nichenet^23^, a computational method that uses prior knowledge on signaling and gene expression networks to predict ligand-target relationships of interacting cells based on their expression data. Astrocytes already have known ligands, such as *APOE*, that bind targets on pericytes with downstream signalling changes that influence pericyte function^75^. Thus, we used Nichenet^23^ and our snRNAseq data to discover and prioritize brain vascular targets that are influenced by astrocytic ligands. In order to identify those vascular target-astrocyte ligand pairs that are most perturbed in AD and therefore most likely to influence BBB dysfunction, we restricted our analyses to those genes that are significant DEGs in these cell types in our data.

Our astrocyte ligand-vascular target analysis revealed strong predicted interactions and 24 brain vascular target candidates most of which had biological functions involving signalling, angiogenesis and cytoskeleton structure. We selected 6 predicted vascular target genes representing each functional category for validation of their differential expression in our snRNAseq cohort using an orthogonal gene expression measurement approach of qPCR. All 6 genes (*ANGPT2*, *AHNAK*, *ECE1*, *TSC22D3*, *STAT3*, *SMAD3*) could be validated for their differential expression in AD vs. control nuclei. Additionally, all genes had positive associations with AD-related neuropathologies, consistent with their higher levels in AD brains.

Of the signalling molecules*^61,68,69^* identified in our study, *SMAD3* and *STAT3* also have roles in vascular function^69,75^. Both of these genes were significantly upregulated in AD brains in the activated pericyte cl.25 and had strong interactions with astrocyte ligands, some of which are known AD risk genes, namely *APOE*, *APP*, *PSEN1* and *MAPT* previously shown to lead to BBB dysfunction in model systems^76^. Both *SMAD3* and *STAT3*, which have two of the highest number and strength of astrocyte ligand interactions amongst all vascular targets in our study, are signalling molecules downstream of many ligands including TGF-β and VEGFR2-binding growth-factors, respectively^77^. Crosstalk between SMAD3 and STAT3 has been demonstrated in numerous conditions^61,77^, especially in cancer. Deletion of Stat3 in astrocytes reduced Aβ in the APP/PSEN1 mouse model of amyloidosis^68^. To our knowledge, neither of these signalling molecules with variable functions have been investigated in human AD brains nor for their roles in BBB dysfunction in AD.

Two other vascular targets in our study with known Aß-related functions were *ECE1^66,78^* and *TSC22D3^70^*, both of which were upregulated in activated pericytes. *ECE1* encodes endothelin converting enzyme-1, which is an Aβ degrading enzyme^66^ that also leads to vasoconstriction through conversion of endothelin precursors to endothelin-1^67^ and is involved in regulating monocyte passage through BBB^79^. *TSC22D3*, which encodes glucocorticoid induced leucine zipper (GILZ) in the CNS, is postulated to have beneficial roles in AD by regulating Aß^80^ or its anti-inflammatory functions^63,80^. Interestingly, *ECE1* and *TSC22D3* both have strong predicted astrocytic ligands that are AD risk genes also with Aß-related functions, namely *APP* and *APOE*, respectively^81^. Our findings suggest that astrocytic expression of these AD risk genes may be regulating pericytic expression of *ECE1* and *TSC22D3*, although this requires experimental confirmation in model systems.

Amongst the vascular targets we identified, *AHNAK*, which is upregulated in activated pericyte cl.25, has the second strongest level of astrocyte ligand interactions and one of five targets with cytoskeleton-related functions (*AHNAK*, *DMD*, *SLC38A2*, *STARD13*, *TIMP3*, **Figure. 4A**). *AHNAK* encodes a 700 kDa structural scaffolding protein which is involved in the integrity and permeability of the BBB^57^. AHNAK is overexpressed in response to BBB injury^82^ and is also upregulated by angiogenic growth factor angiopoietin-1^57^. Notably, two angiogenesis genes, *ANGPT2^65^* and *AMOTL2^64^* were also vascular targets in our study and were upregulated in the endothelial cluster cl.27. *ANGPT2*, the most significantly upregulated vascular target gene in AD nuclei in this cluster, is a growth factor with signalling properties^65^, like the strong activated pericyte cl.25 targets *STAT3* and *SMAD3*.

Dysfunctions in cellular interactions and signaling in the GVU is critical to understand the mechanisms underlying BBB dysfunction that contributes to AD pathophysiology^74,76^. Our study demonstrates transcriptional alterations of vascular cells and astrocytes of GVU in AD at single cell resolution and discovers target-ligand relationships between these cell types. These findings are expected to pave the way to uncover novel mechanistic interactions between pericytes, endothelia and astrocytes and their perturbations in AD. Despite these strengths, our study also has some weaknesses. In this study we focused on predicted interactions of brain vascular target molecules with astrocytic ligands, given their known crosstalk at the BBB^11,75^, however it will be important to also interrogate interactions with neurons. Although we have validated the expression changes in AD of select vascular targets with the orthogonal qPCR approach, the interactions *per se* also need to be validated by detailed cell biology approaches that are beyond the scope of our study. Finally, given that there are no snRNAseq studies of pericytes to our knowledge, our findings for this rare cell type needs to be validated in future studies.

In summary, using snRNAseq in AD and elderly control temporal cortex tissue, we evaluated 79,751 high quality nuclei in 35 clusters, of which 4,604 nuclei formed three distinct brain vascular cell clusters of activated pericytes (cl.25), endothelia (cl.27) and resting pericytes (cl.31). AD brains have significant DEGs in all three clusters many of which are cluster-specific and most of which reside in activated pericytes, demonstrating widespread transcriptome perturbations in this important cell type of the BBB. We uncovered computationally predicted interactions between astrocytic ligands and vascular targets, which underscores potential downstream effects of transcriptional changes at the GVU. We identified target-ligand interactions for genes that are both well-known for AD risk such as *ECE1-APP*, and those that are novel. Collectively, our study provides a prioritized list of perturbed brain vascular molecules, their astrocytic partners at the GVU in AD and offers mechanistic avenues to explore for deciphering the precise molecular mechanisms of BBB dysfunction in AD.

## Abbreviations

Aß: Amyloid ß
AD: Alzheimer’s disease
BBB: Blood-brain barrier
CNS: Central nervous system
DEG: Differentially expressed genes
ECM: Extracellular matrix
FACS: Fluorescence-activated cell sorting
FANS: Fluorescence-activated nuclear sorting
GO: Gene ontology
GVU: Gliovascular unit
QC: Quality control
RNAseq: RNA sequencing
scRNAseq: Single cell RNA sequencing
snRNAseq: Single nucleus RNA sequencing
TCX: Temporal cortex
UMAP: Uniform Manifold Approximation and Projection
UMI: Unique molecular identifier

## Methods

### Brain donors and samples

Frozen post-mortem brain tissues from 12 AD patients and 12 control donors, matched for age at death and sex, were obtained from the Mayo Clinic Brain Bank. Total RNA from ~20 mg collected tissue was isolated to evaluate tissue quality. RNA integrity number (RIN) was determined via Agilent 2100 Bioanalyzer using RNA Pico Chip assay and tissues that have RIN > 6.0 were utilized in nuclei isolation and single nucleus RNA sequencing (snRNAseq).

### Histology and immuno-histochemistry

Neuropathological assessment that comprises evaluation of gross and microscopic findings, as well as quantitative analysis of Alzheimer type pathology was conducted. Braak neurofibrillary tangle (NFT) and Thal amyloid stages were assigned as previously described^83^. Presence of TDP43 inclusion bodies were determined by immunohistochemistry with antibodies directed against pathological TDP43^84^.

### Nuclei isolation

Single nuclei suspensions were collected from human temporal cortex. 100 mg tissue was directly transferred from dry ice to dounce homogenizer that contains homogenization buffer (0.25 M sucrose, 25 mM KCl, 5 mM MgCl2, 20 mM tricine-KOH, pH 7.8, 1 mM DTT, 0.15 mM spermine, 0.5 mM spermidine, protease inhibitors, 5 μ g/mL actinomycin, 5 u/ μL recombinant RNAase inhibitor, and 0.04% BSA). Twenty-five strokes with loose and tight pestle were performed, sequentially. After strokes with tight pestle, 5% IGEPAL (Sigma) solution was added to reach a final concentration of 0.32%. Ten additional strokes were performed, and homogenate was filtered through 30 μm cell strainers. Filtrated homogenate was centrifuged down (500g, 5 minutes), and washed once with wash and storage buffer (1XPBS with 2%BSA and 5 u/ μL recombinant RNAase inhibitor). After washing, homogenate was filtered again through 30 μm cell strainer and centrifuged down for 10 minutes at 500g. The pellet was re-suspended in 700μl cold PBS with 5 U/μl RNAse inhibitors. 300μl debris removal solution (Miltenyi Biotech) was added and the solution was gently mixed. The solution was carefully overlaid with 1 mL wash and storage buffer (WSB) and centrifuged for 10 minutes at 3000g. Supernatant was removed, and the pellet was washed with WSB and centrifuged for 10 minutes at 1000g.

### Flow cytometry and nuclei sorting (FANS)

Isolated nuclei were incubated with Human Nuclear Antigen [235-1] (ab191181, Abcam) antibody at 1:200 for 1 hour on ice. Mouse IgG1, kappa monoclonal isotype control was included in the staining. Nuclei were incubated for 30 minutes on ice in secondary antibody solution that contains goat anti mouse 488 secondary antibody. Nuclei were then resuspended and sorted into WSB via BD FACSAria II sorter.

### Quality control of isolated nuclei

To assess the purity of the sorted nuclei, both RNA and protein profiles were analyzed. RNA was isolated via Qiagen In RNA level, disappearances of 18S and 28S rRNA peaks in Agilent histogram were analyzed to confirm lack of cytoplasmic RNA contaminants in the nuclei preparations. Nuclear H3 and mitochondrial COX4 protein ratios were checked via western blot to confirm nuclear purity in protein level. Also, to confirm the preparation method does not cause bias in favor of certain cell type, qPCR was performed with probes against RNU2.1 (Nuclear probe), AQP4, CD34, P2RY2, RBFOX3, and MOG. Nuclei integrity was checked under microscope with 20X objective of Evos Cell Imaging System and through Z-stah images captured with 100X objective of Confocal Laser Scanning Microscope.

### 10X cDNA library production and snRNAseq

To quantify the number of sorted nuclei, nuclei was stained with 0.04 % trypan blue and counted in a hemocytometer. Total nuclei solution was diluted to 1000 nuclei/μl. A total of 3000 estimated nuclei were loaded on the 10X Chromium microchip and run on a Single Cell Instrument (10X Genomics) to generate single cell gel beads-in-emulsion (GEMs). Single cell RNAseq libraries were prepared using the Chromium Single Cell 3’ Gel Bead and Library Kit v3 (10X Genomics, No. 120237) and the Chromium i7 Multiplex Kit (10X Genomics, No. 120262) according to the manufacturer’s instructions. Qualities of libraries were checked using Agilent High Sensitivity DNA Kit via Agilent 2100 Bioanalyzer.

DNA libraries were sequenced by the Mayo Clinic Medical Genome Facility (MGF) core to obtain expression measures on the Illumina HiSeq4000 sequencer. Two samples were run on each lane of one flow cell and two flow cells were used in total.

### Reads alignment and quality control

10X Genomics Cell Ranger Single Cell Software Suite v3.1.0 were used to demultiplex raw base call files generated from the sequencer into FASTQ files. Raw reads were aligned to human genome build GRCh38 and a premature mRNA reference file. Reads aligned to gene transcript locus, including both exonic and intronic regions, were counted to generate raw UMI counts per gene per barcode for each sample. The raw UMI matrices were filtered to only keep barcodes with ≥ 500 UMIs and those that were called a ‘cell’ by Cell Ranger’s cell calling algorithm. The filtered barcodes from all 24 samples were pooled together and further filtering criteria were applied to exclude the following barcodes and genes. 1) barcodes with > 10% of UMI mapped to mitochondrial genome; 2) barcodes with < 400 or > 8000 detected genes; 3) barcodes with < 500 or > 46425 mapped UMIs; 5) genes that are detected in < 5 cells. The above thresholds were determined by UMI or gene distribution to identify undetectable genes and outlier barcodes that may encode background, broken or multiple cells. Next, we extracted protein coding genes for further analysis. Recorded sex of samples was compared to the sex inferred from chromosome Y gene expression, which confirmed the correctness of sex information of those samples.

### Clustering nuclei

After quality control, UMI counts of remaining cells and genes were normalized using NormalizeData function in R package Seurat^85^ v3.1.0, which gave natural log transformed expression adjusted for total UMI counts in each cell. The top 2000 genes whose normalized expression varied the most across cells were identified through FindVariableFeatures function with default parameters. Using those genes, cells from eight groups of samples (grouped by AD/normal, male/female and *APOEε4* positive/negative) were integrated using functions FindIntegrationAnchors and IntegrateData with default parameters. Principal components (PCs) of the integrated and scaled data were computed; and the first 31 PCs, which accounted for > 95% variance, were used in clustering cells. Cell clustering was performed using FindNeighbors and FindClusters with default parameters. All analyses described this section were performed using Seurat v3.1.0.

### Identifying cluster marker genes and assigning cell types of each cluster

Marker genes that were conserved in both AD and control nuclei were identified in each cluster using FindConservedMarkers in Seurat v3.1.0. Marker genes of one cluster must 1) be presented in > 20% AD nuclei and > 20% control nuclei of this cluster; 2) the log(fold change) between their expression in AD (control) cells of this cluster and AD (control) cells of other clusters must be >0.25; 3) the rank sum test p-value (Bonferroni adjusted) between AD (control) cells in this cluster and AD (control) cells in other clusters < 0.05.

Two approaches were adopted and combined for cell type assignment. The first one utilized the marker gene lists reported in R BRETIGEA^26^ for neurons (1000 markers), astrocytes (1000 markers), oligodendrocytes (1000 markers), microglia (1000 markers), endothelial cell (1000 markers) and OPCs (500 markers). Hypergeometric tests were performed for over-representation of our cluster markers in those reported markers. Each cluster was assigned one cell type that was most over-represented. The second approach was to check the existence of a handful of well-recognized cell type markers in top cluster markers. Those cell type markers are *SYT1*, *GRIN1* for neuron; *SLC17A7*, *NRGN* for excitatory neuron; *GAD1*, *GAD2* for inhibitory neuron; *VCAN*, *PDGFRA* for OPC, *MBP*, *MOBP* for oligodendrocyte; *C3*, *CD74* for microglia; AQP4, GFAP for astrocyte; *FLT1*, *PECAM1/CD31* for endothelial cells; and *PDGFRB*, *PDE5A* for pericytes. Combining the two approaches, we assigned the following eight cell types/subtypes to each cluster - excitatory neuron, inhibitory neuron, oligodendrocyte, OPC, microglia, astrocyte, endothelia and pericytes.

### Cell distribution association test

For each cluster, the number of cells was divided by the total number of cells in all clusters for a given individual. The resulting ratio gives the cell distribution that was used to test for association with characteristics using a Wilcoxon rank sum test for binary variables (AD vs. control, male vs. female, *APOEε4* positive vs. negative, and TDP43 positive vs. negative) or Spearman’s test of correlation for quantitative/semi-quantitative variables (age at death, Thal phase, and Braak stage). All statistical tests were two-sided. P-values <0.05 were considered statistically significant.

### Differential expression and association analysis for each cluster

For each cluster, we performed differential expression analysis for genes that were detected (UMI >= 1) in >= 10% AD cells or >= 10% normal cells using R package MAST ^53^. MAST employed a hurdle model to accommodate the so-called 0-inflation observed in scRNAseq/snRNAseq data, i.e., many cells had 0 UMI for a given gene. In the following models, AD cells were coded as 1, normal cells were coded as 0; males were coded as 1, females were coded as 0; APOE4 positive (44 or 24 or 34) were coded as 1, APOE4 negative were coded as 0; TDP43 positive were coded as 1, TDP43 negative were coded as 0; age, Braak and Thal stages were numerical variables.

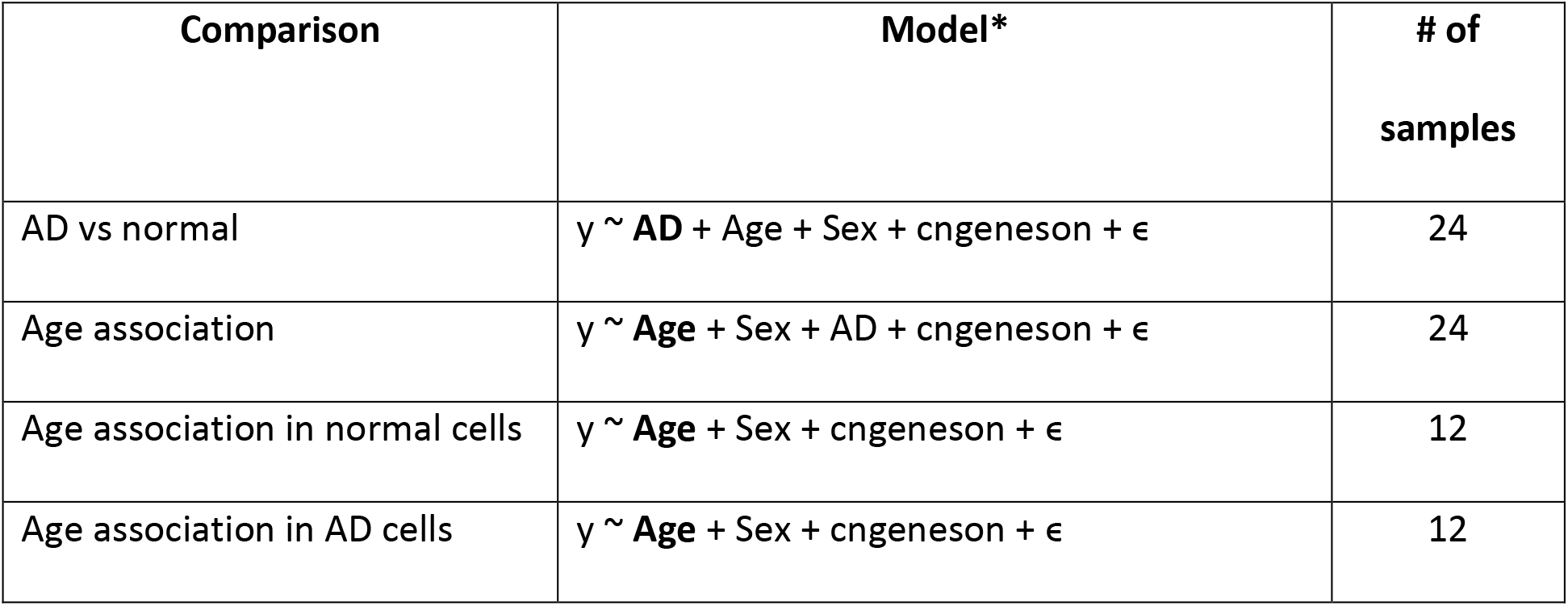
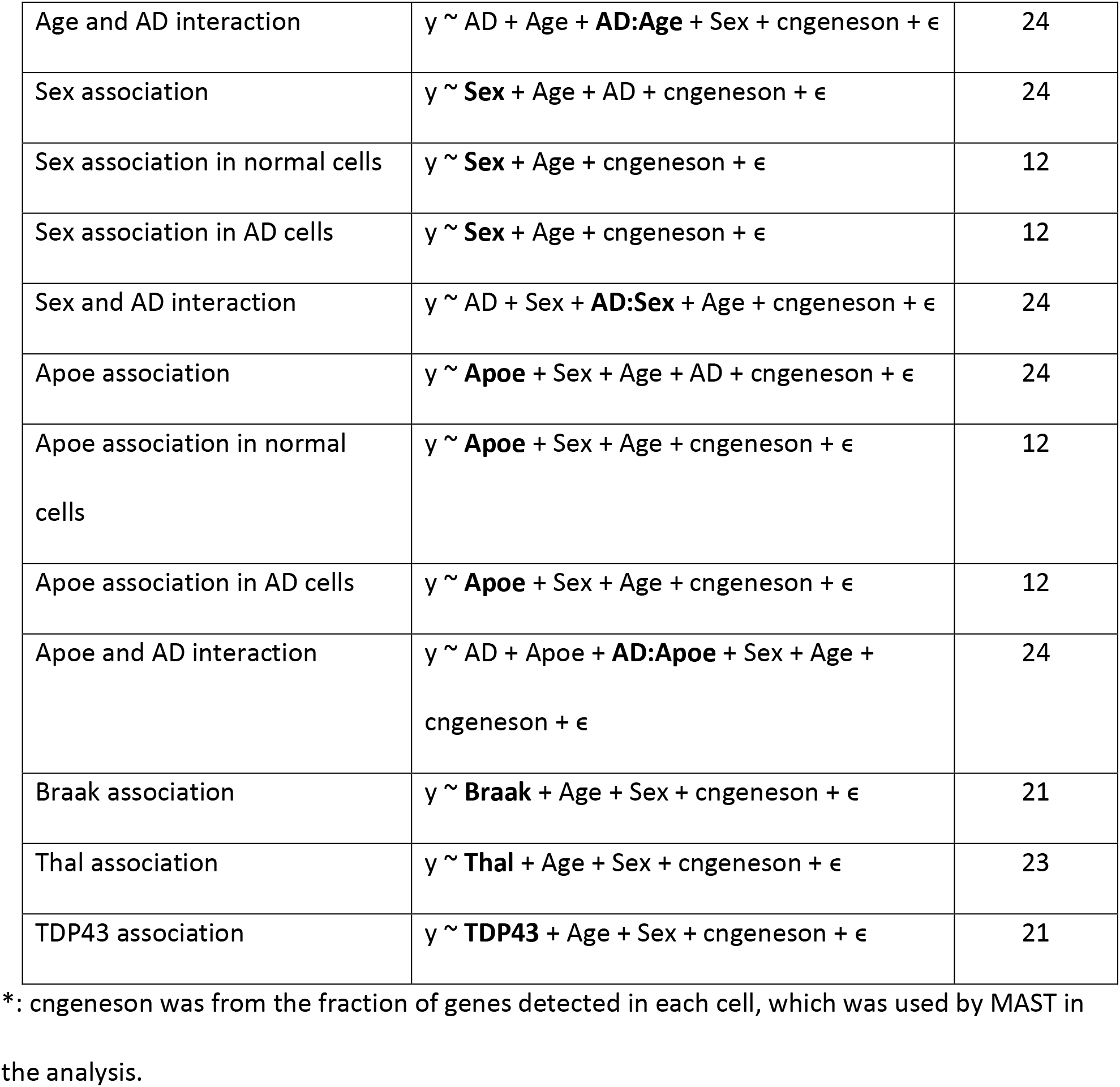

| Comparison | Model* | # of samples |
| --- | --- | --- |
| AD vs normal | $y \sim \mathbf{AD} + \text{Age} + \text{Sex} + \text{cngeneson} + \epsilon$ | 24 |
| Age association | $y \sim \mathbf{Age} + \text{Sex} + \text{AD} + \text{cngeneson} + \epsilon$ | 24 |
| Age association in normal cells | $y \sim \mathbf{Age} + \text{Sex} + \text{cngeneson} + \epsilon$ | 12 |
| Age association in AD cells | $y \sim \mathbf{Age} + \text{Sex} + \text{cngeneson} + \epsilon$ | 12 |
| Age and AD interaction | $y \sim \text{AD} + \text{Age} + \mathbf{AD:Age} + \text{Sex} + \text{cngeneson} + \epsilon$ | 24 |
| Sex association | $y \sim \mathbf{Sex} + \text{Age} + \text{AD} + \text{cngeneson} + \epsilon$ | 24 |
| Sex association in normal cells | $y \sim \mathbf{Sex} + \text{Age} + \text{cngeneson} + \epsilon$ | 12 |
| Sex association in AD cells | $y \sim \mathbf{Sex} + \text{Age} + \text{cngeneson} + \epsilon$ | 12 |
| Sex and AD interaction | $y \sim \text{AD} + \text{Sex} + \mathbf{AD:Sex} + \text{Age} + \text{cngeneson} + \epsilon$ | 24 |
| Apoe association | $y \sim \mathbf{Apoe} + \text{Sex} + \text{Age} + \text{AD} + \text{cngeneson} + \epsilon$ | 24 |
| Apoe association in normal cells | $y \sim \mathbf{Apoe} + \text{Sex} + \text{Age} + \text{cngeneson} + \epsilon$ | 12 |
| Apoe association in AD cells | $y \sim \mathbf{Apoe} + \text{Sex} + \text{Age} + \text{cngeneson} + \epsilon$ | 12 |
| Apoe and AD interaction | $y \sim \text{AD} + \text{Apoe} + \mathbf{AD:Apoe} + \text{Sex} + \text{Age} + \text{cngeneson} + \epsilon$ | 24 |
| Braak association | $y \sim \mathbf{Braak} + \text{Age} + \text{Sex} + \text{cngeneson} + \epsilon$ | 21 |
| Thal association | $y \sim \mathbf{Thal} + \text{Age} + \text{Sex} + \text{cngeneson} + \epsilon$ | 23 |
| TDP43 association | $y \sim \mathbf{TDP43} + \text{Age} + \text{Sex} + \text{cngeneson} + \epsilon$ | 21 |

\*: cngeneson was from the fraction of genes detected in each cell, which was used by MAST in the analysis.

For selecting DEGs or genes associated with continuous variables from each cell cluster, we focused on the set of genes that were detected (UMI>0) in at least 20% of the AD cells or of the normal cells in the cluster and have q < 0.05. The DEGs between binary variable (including AD and normal, male and female, APOE4 positive and APOE4 negative, or TDP-43 yes and TDP-43 no) for each cluster were the ones that have q < 0.05 and |logFC| > 0.1.

### Signature genes of clusters

Signature genes of a cluster are genes that highly expressed in one cluster of a cell type but not the other clusters of that cell type such that a) they are present in >= 50% cells of this cluster, b) average log2 fold change >= 1.0 and c) Wilcoxon rank sum test Bonferroni p-value < 0.05 when compared to each of the other other clusters. FindMarker function of Seurat was applied to obtain such signature genes. Signature genes for vascular and astrocyte clusters were both determined as above.

### Pseudotime analysis of clusters

Pseudotime analysis was performed using monocle3 R package as follows. The principal graph was learned using multi tree DDRTree method from the UMAP coordinates of existing clusters, which were obtained from Seurat utilizing the top 2000 most variable genes as described previously. We then set the first principal graph node in one of the clusters (cluster 25 for vascular and cluster 27 for astrocyte) as root state. Pseudotime was calculated between each cell and the root state.

### Enrichment of genes in MSigDB GO terms

MSigDB v7.0 was used for these analyses. The enrichment of selected genes in MSigDB C5 category (i.e., gene ontology or GO) was performed using R enRichment package. The top 5 enriched GO biological process (BP) terms and top 5 genes that occur most frequently in these terms were plotted for the main figures.

### Constellation plot

This analysis used script from Olah et al. ^7^ at github.com/vilasmenon/Microglia_Olah_et_al_2020. For every pair of clusters, cells from the two clusters were randomly divided into four groups. For each group, cells in the other three group were used as training data and the cells of this group were classified to be from one of the two clusters. This classification procedure was repeated 100 times and therefore each cell was classified 100 times. If a cell was misclassified > 25 times, it was considered as “ambiguous” or “intermediate”. The percent of intermediate cells was calculated as 100* (num of intermediate cells)/(num.cell.clusterA + num.cell.clusterB).

### Ligand-target analyses

NicheNet^23^ analysis tool was used to study the interaction between astrocytic and vascular cells through nichenetr R package. Prior knowledge of ligand-target interaction has been compiled and optimized by NicheNet from multiple data sources to give a prior model which contains the regulation strength of ligands towards target genes. In this study, we set astrocyte DEGs between AD and normal cells from cl.9, cl.12 and cl.26 as potential ligands, and DEGs from vascular clusters cl.25, cl.27 and cl.31 as target genes. The following description uses activated pericyte cl.25 as an example. Among the potential ligands, we identified genes satisfying the following: a) It is a ligand according to prior model; b) It has receptor genes expressed in the target cluster pericyte cl.25. For the resulting set of ligands, we identify the target genes satisfying that a) It is a DEG of activated pericyte cl.25 and b) It is among the top 250 regulated genes by one of the ligands according to prior model.

In this manner, we obtained cl.9-cl.25, cl.12-cl.25 and cl.26-cl.25 targets interacting with ligands in the aforementioned astrocyte cluster ligands. The intersection of these three sets of targets gives 19 target genes in cl.25. Using a similar approach we identified 6 target genes from cl.27 and 1 target gene from cl.31.

### Gene Expression Validation via qPCR

Total RNA was extracted from sorted nuclei using the miRNeasy Serum/Plasma Kit (QIAGEN; 217184). The Agilent BioAnalyzer RNA 6000 Pico Kit (Agilent; 5067-1514) was used to assess RNA concentration and quality. RNA was normalized to 0.5ng/μl for cDNA synthesis using the SuperScript IV VILO Master Mix (ThermoFisher; 11756050). TaqMan PreAmp Master Mix (ThermoFisher; 4391128) was used to pre-amplify cDNA, followed by TaqMan Universal PCR Master Mix (ThermoFisher; 4304437) with the following gene expression probes: MOG, AQP4, RBFOX3, P2RY12, CD34, ANGPT2, AHNAK, ECE1, SMAD3, STAT3, TSC22D3, GAPDH, RNU2-1 (ThermoFisher; Hs01555268_m1, Hs00242342_m1, Hs01370654_m1, Hs00224470_m1, Hs00375822_m1, Hs00169867_m1, Hs01043735_m1, Hs00969210_m1, Hs00374280_m1, Hs00608272_m1, Hs99999905_m1, Hs03023892_g1). RT-qPCR was performed on a QuantStudio 7 Flex Real-Time PCR System (ThermoFisher). Comparative CT analysis (ΔΔCT) was used to quantify gene expression with RNU2-1 used as the endogenous reference and brain homogenate as the calibrator.

