## Supplementary Figures for "Single Nuclei Transcriptome Reveals Perturbed Brain Vascular Molecules in Alzheimer’s Disease"

**Figure S1:**

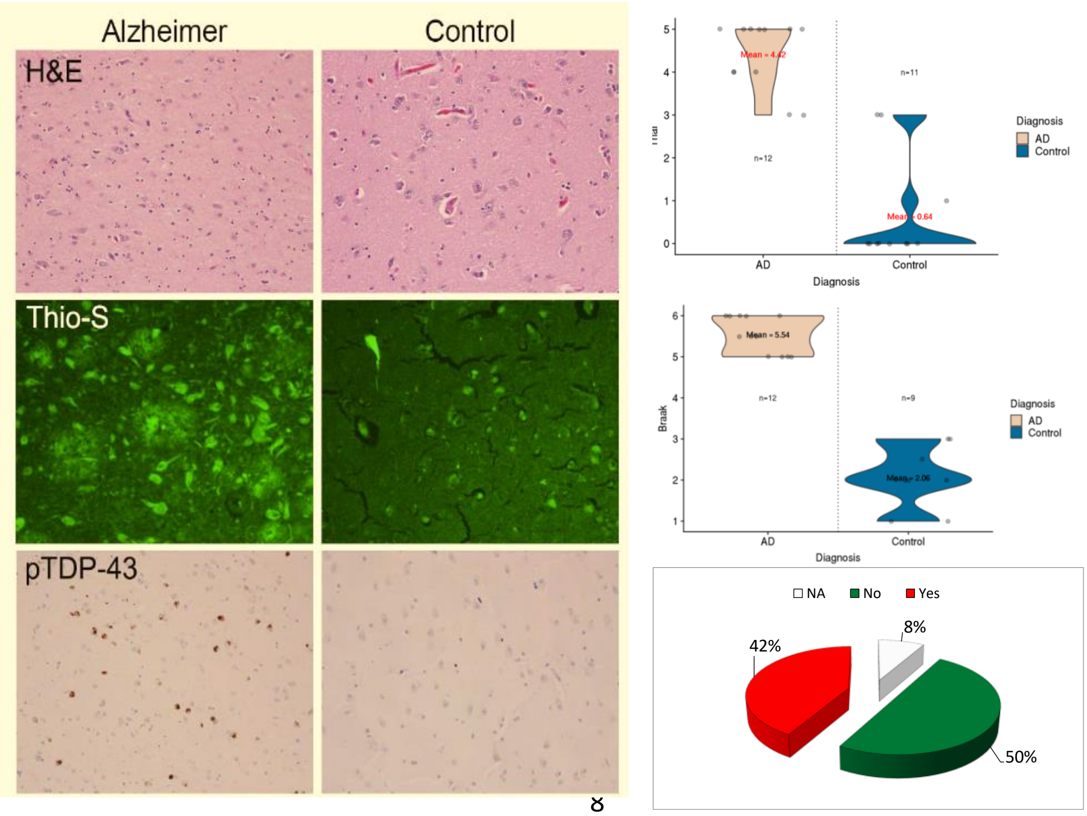


**Figure S1: Neuropathological measures in AD and control samples.** Left panels depict representative immunohistochemistry pictures from an Alzheimer's patient and control brain for H&E, thioflavin-S and TDP-43 staining. Right sided panels from top to bottom show distribution of Thal phase, Braak stage and presence or absence of TDP-43 in the cohort of 12 AD and 12 control brains.

**Figure S2:**

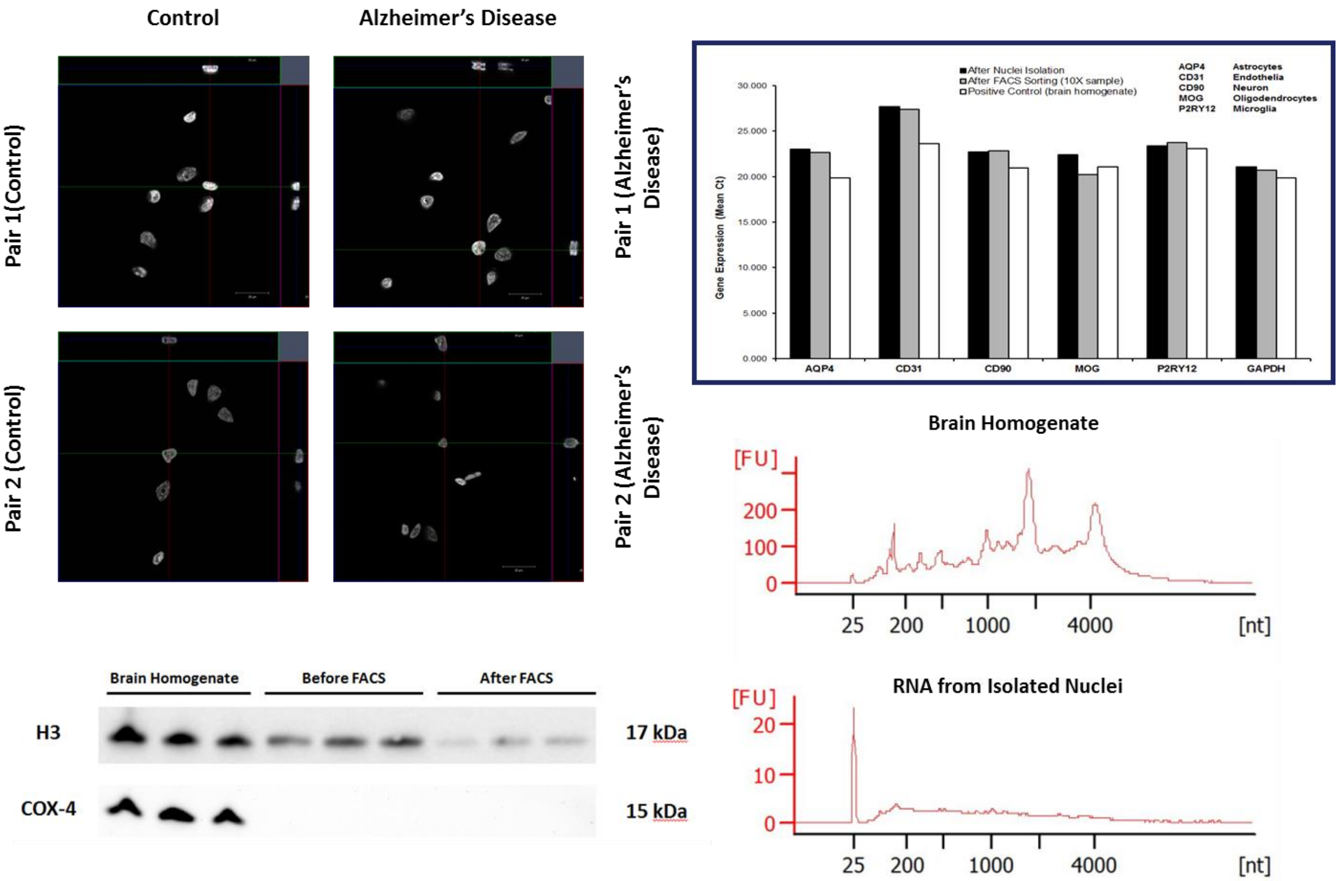


**Figure S2: Purity and quality of isolated and purified nuclei.** Top left: Intact nuclei were observed under confocal microscope. Bottom left: Effect of FANS to purity of isolated nuclei was measured using Nuclear (Anti-H3) and mitochondrial (Anti-COX4) antibodies. Top right: qPCR was performed with cell type specific TaqMan probes from isolated nuclear RNA demonstrate lack of bias in detecting various cell types. Bottom right: Disappearance of 18S and 28S peaks in Bioanalyzer histogram demonstrates the disappearance of cytoplasmic RNA from the nuclear fraction.

**Figure S3:**

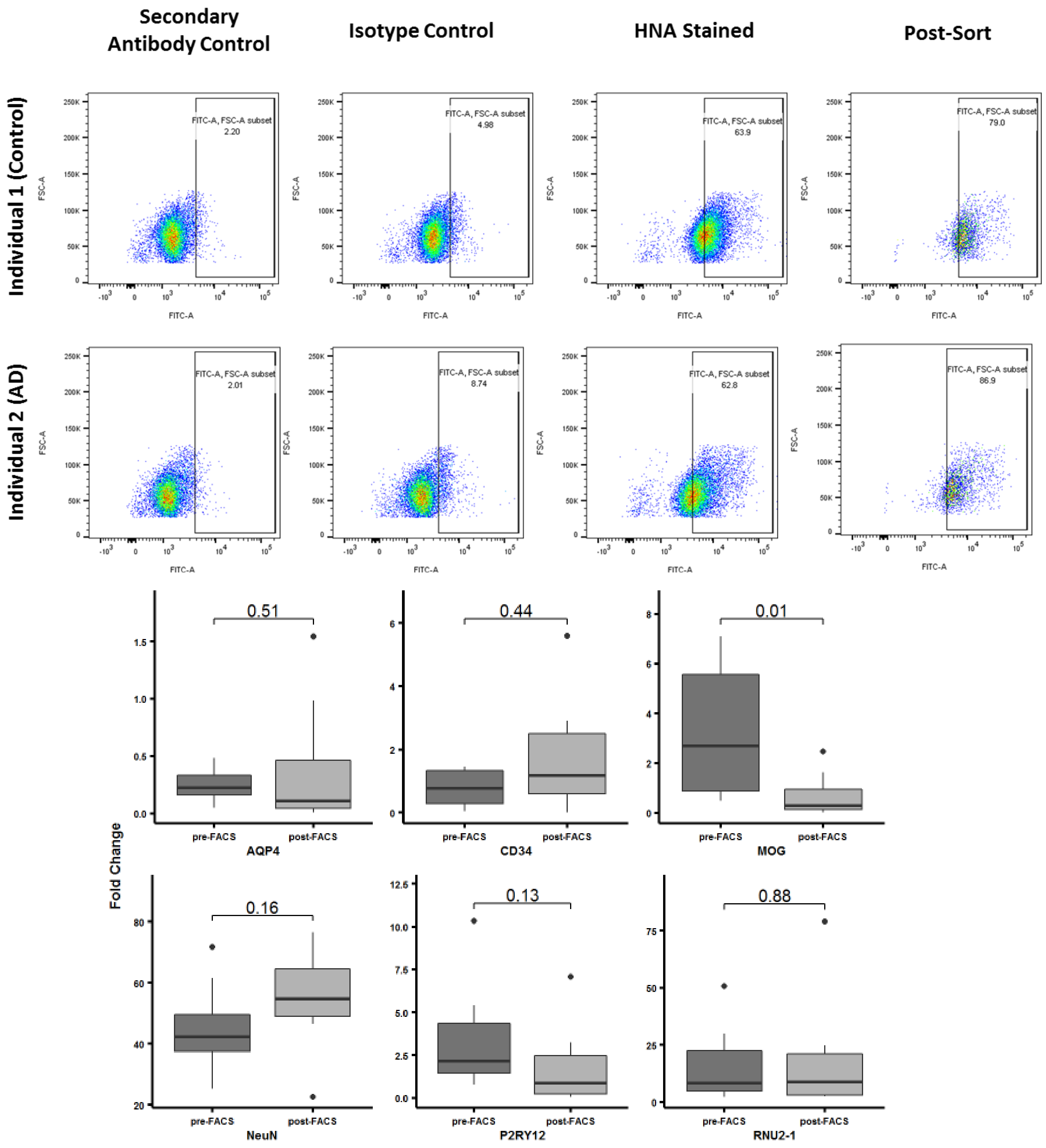


**Figure S3: Fluorescence-activated cell sorting results.** Upper: FACS profile of the sorted nuclei. Lower: effect of FACS sorting to the distribution of cell types.

**Figure S4:**

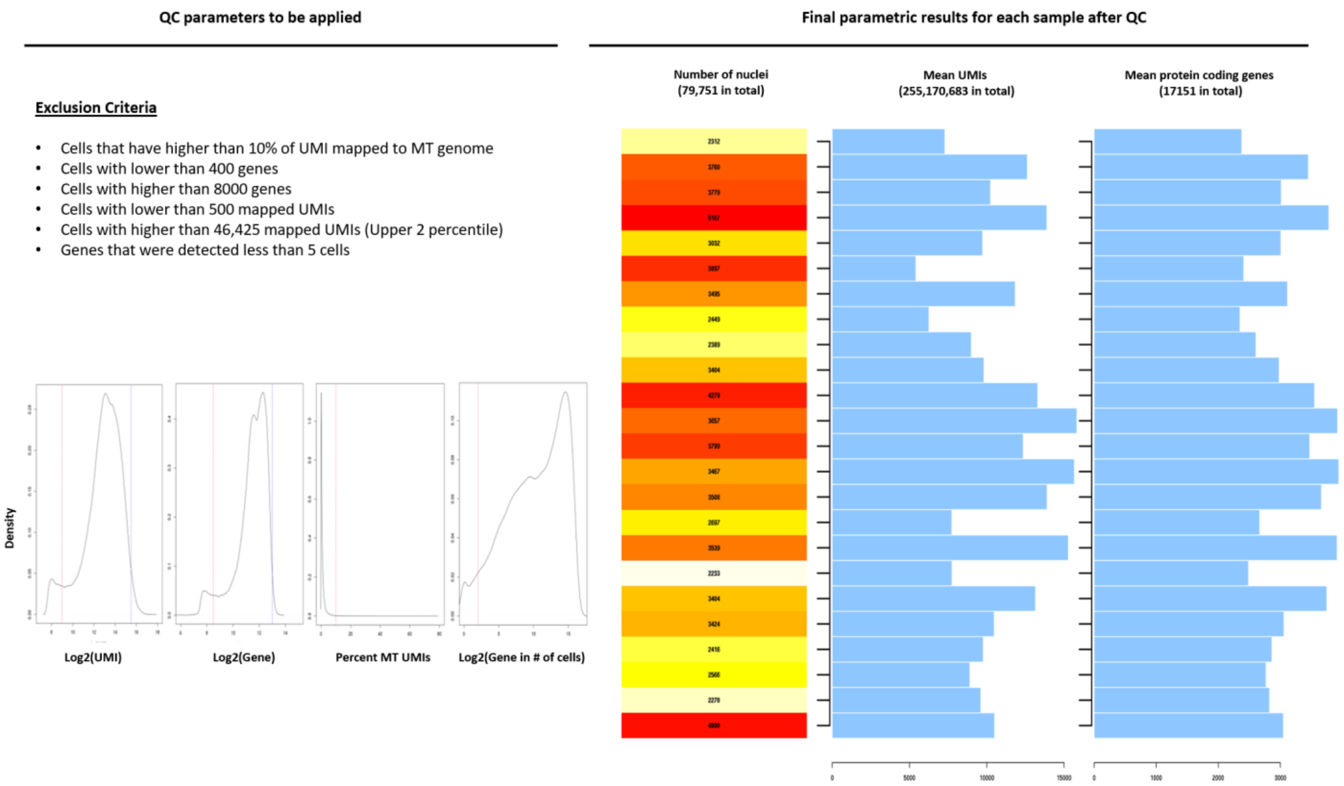


**Figure S4: Parametric results of snRNAseq data after quality control and filtration steps.** Top left: Exclusion criteria for nuclei and genes. Bottom left: Density plots: UMI=unique molecular identifier. Right panel: Post-QC number of nuclei, mean UMI and number of protein coding genes per each sample.

**Figure S5:**

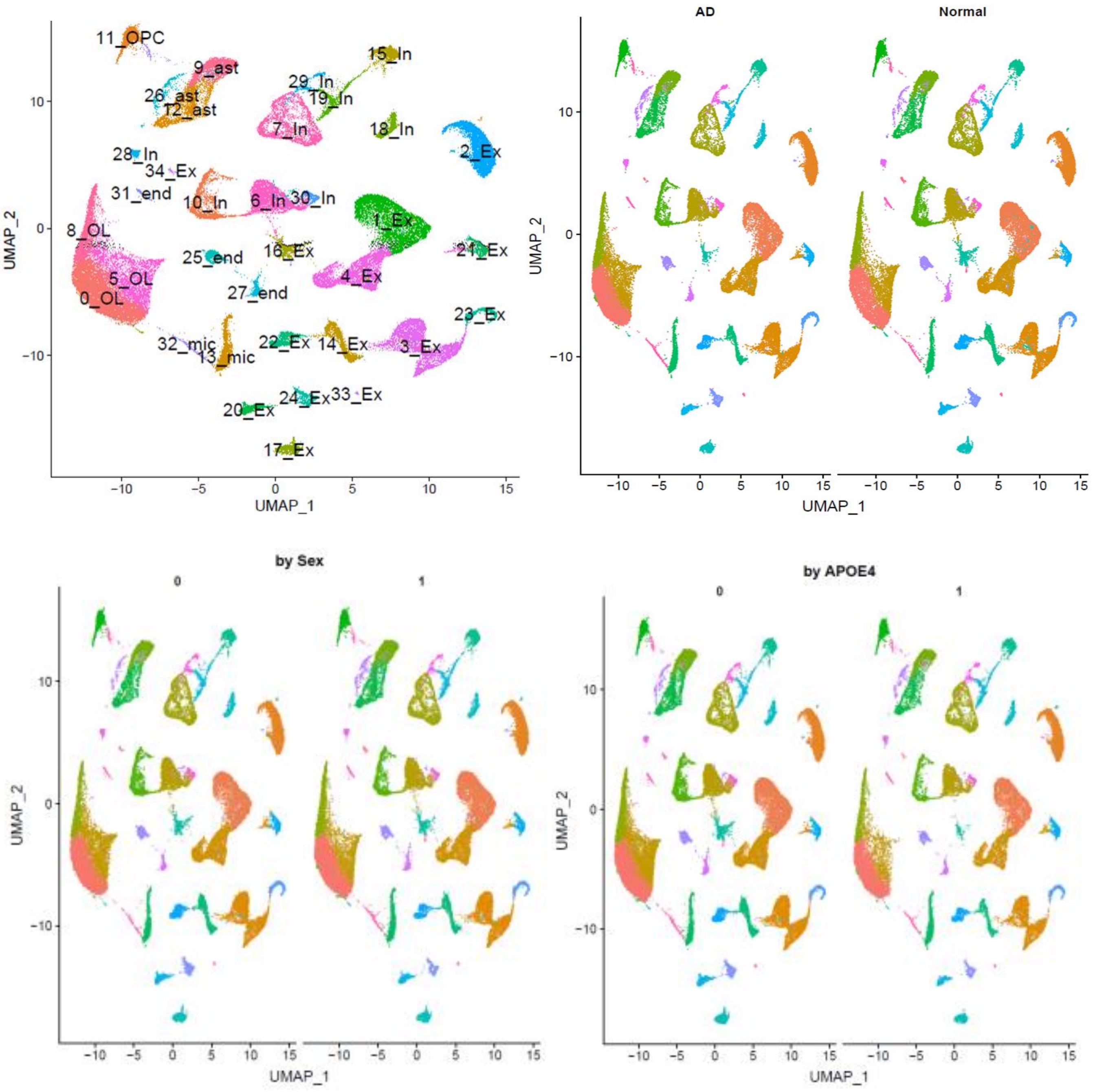


**Figure S5:** Top left: Nuclei clusters displayed on UMAP reduced dimension space, annotated as cluster number followed by cell type. Ex: excitatory neuron; In: inhibitory neuron; OL: oligodendrocyte; OPC: oligodendrocyte progenitor cell; end: endothelia; per: pericyte; ast: astrocyte; mic: microglia. Distributions are also shown stratified by diagnosis (top right); sex (bottom left, 0=female, 1=male) and *APOE* (bottom right, 0=*APOE* ε4 negative, 1=*APOE* ε4 positive).

**Figure S6:**

### A. Cluster nuclei proportions by diagnosis

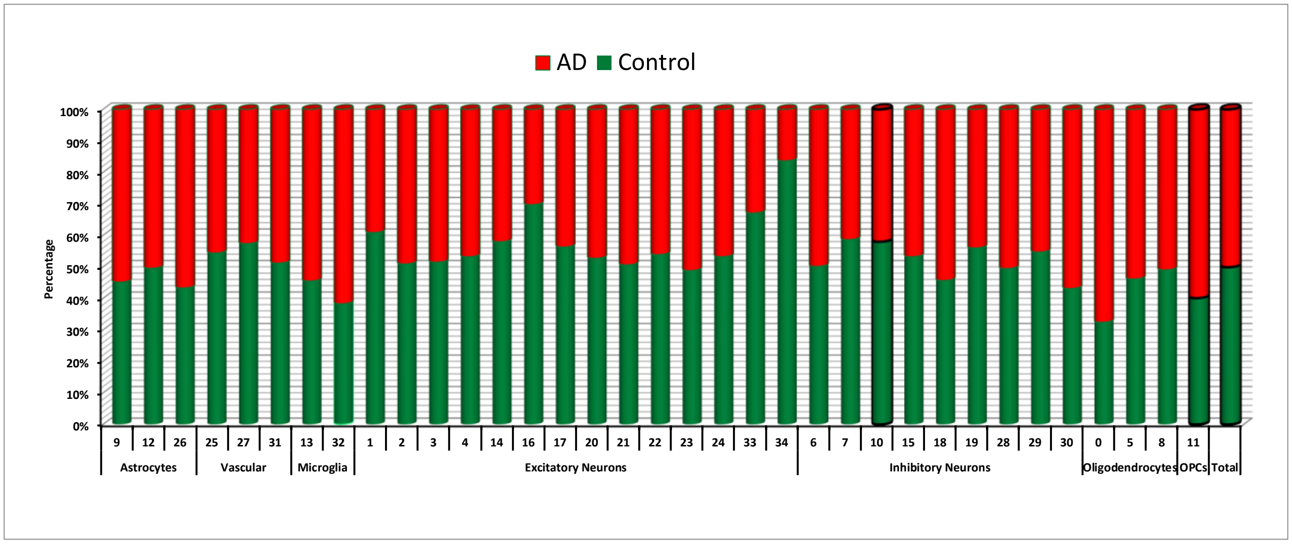


### B. Cluster nuclei proportions by Braak stage

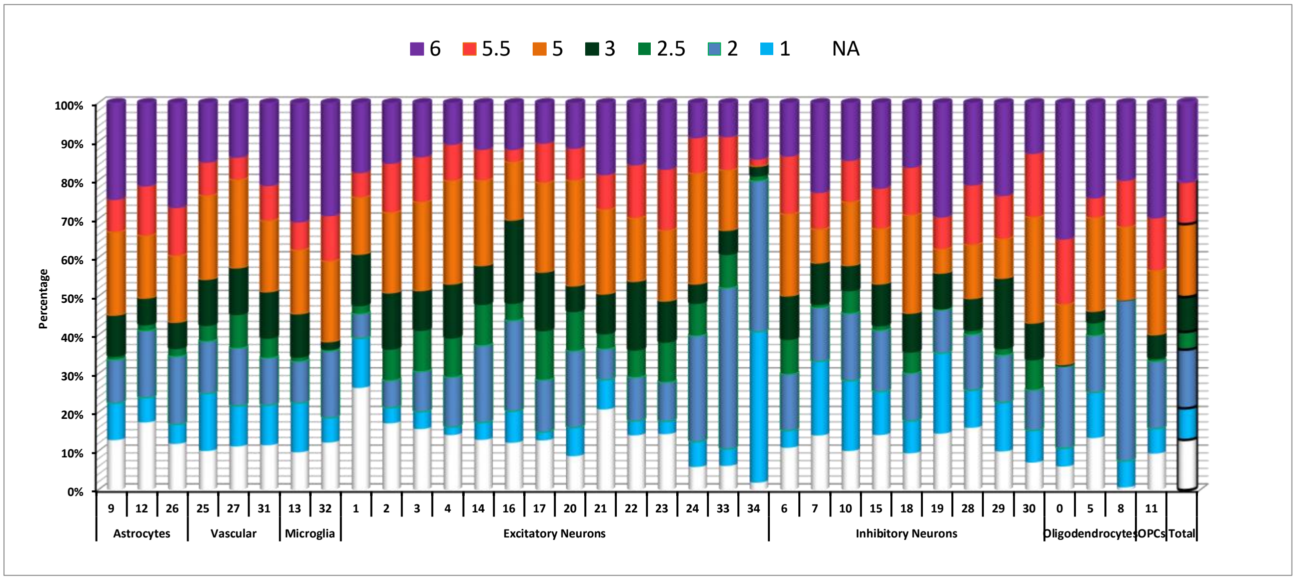

### C. Cluster nuclei proportions by Thal phase

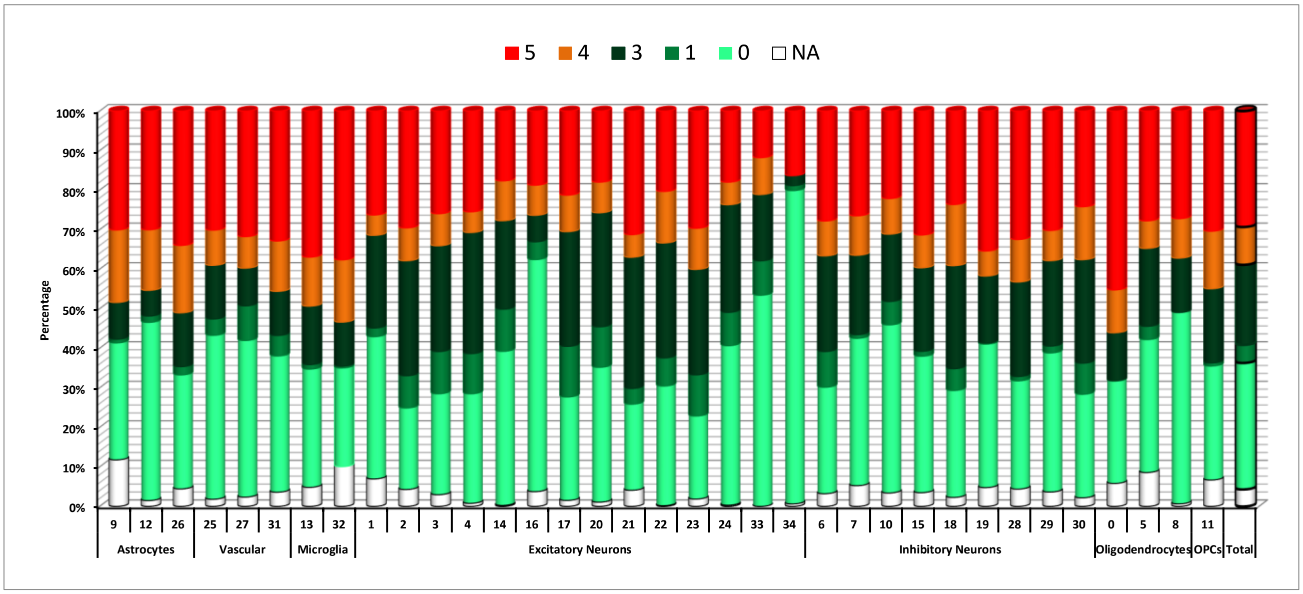


### D. Cluster nuclei proportions by TDP43

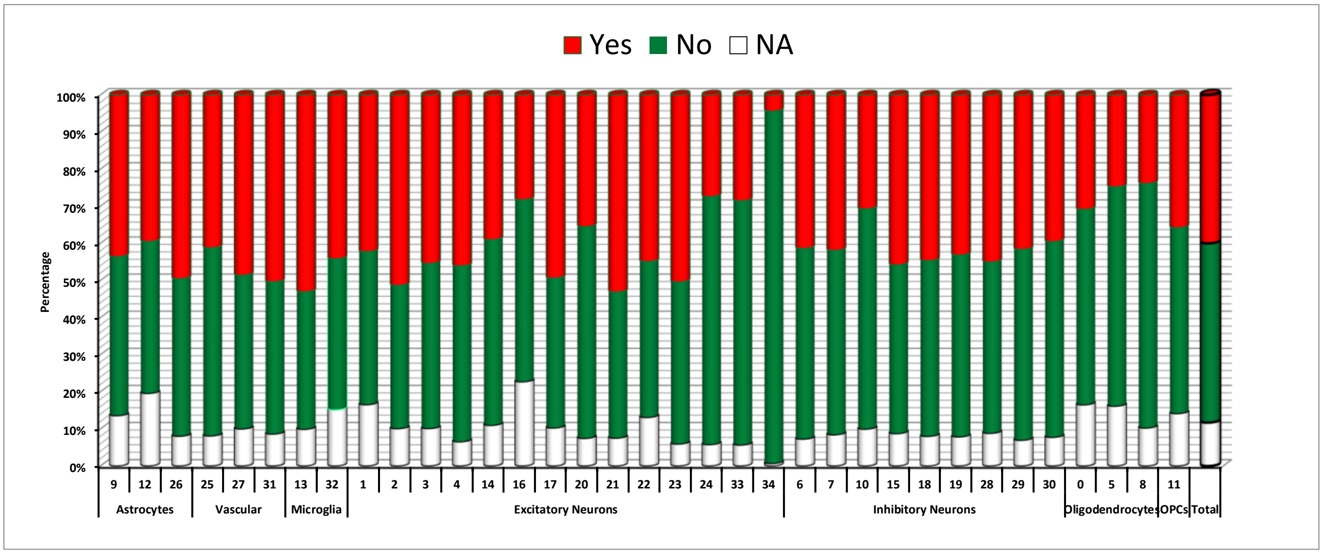

### E. Cluster nuclei proportions by sex

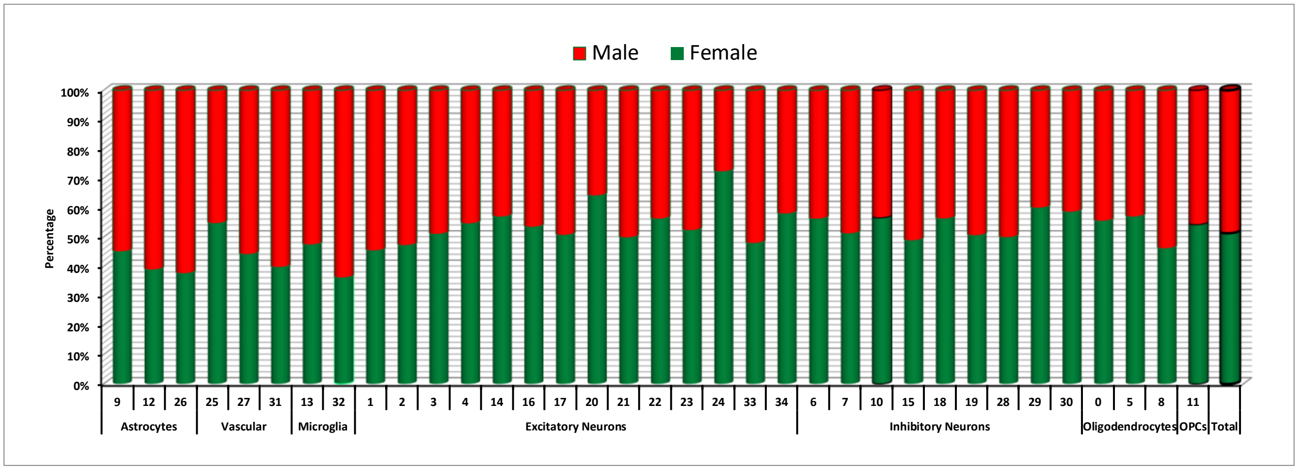


### F. Cluster nuclei proportions by *APOE*

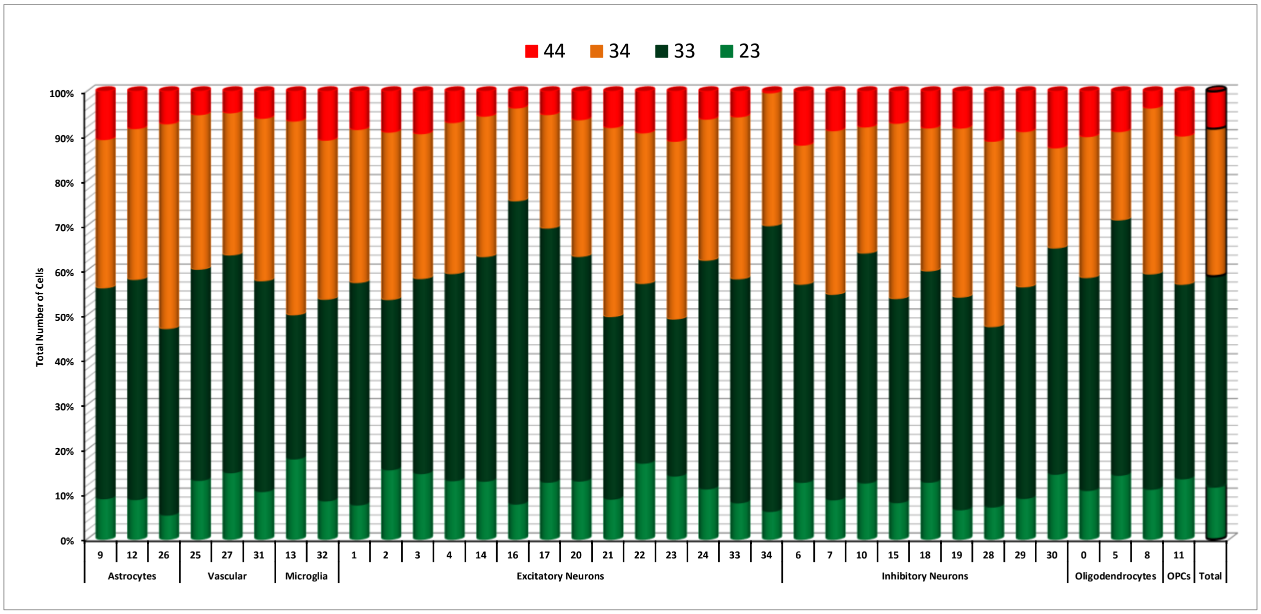

### G. Cluster nuclei proportions by age

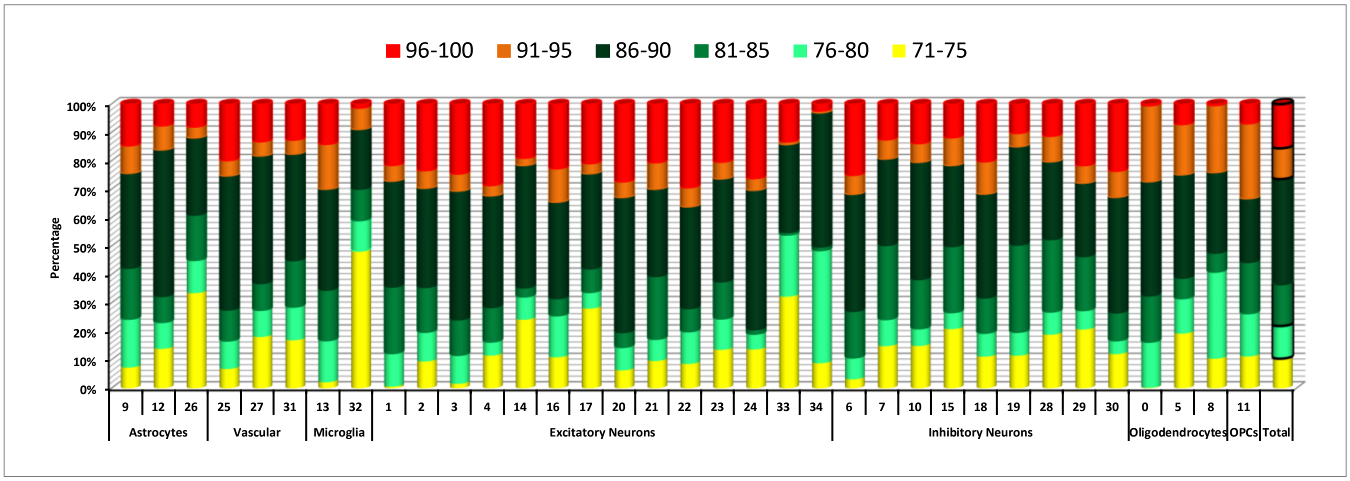


**Figure S6:** Cluster nuclei proportion distributions by a. diagnosis, b. Braak Stage, c. Thal phase, d. TDP-43, e. sex, f. *APOE* and g. age.

**Figure S7:**

**A.**

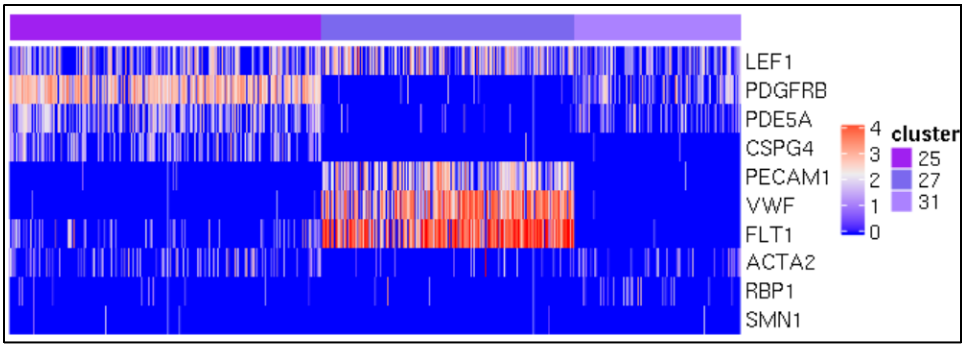


**B.**

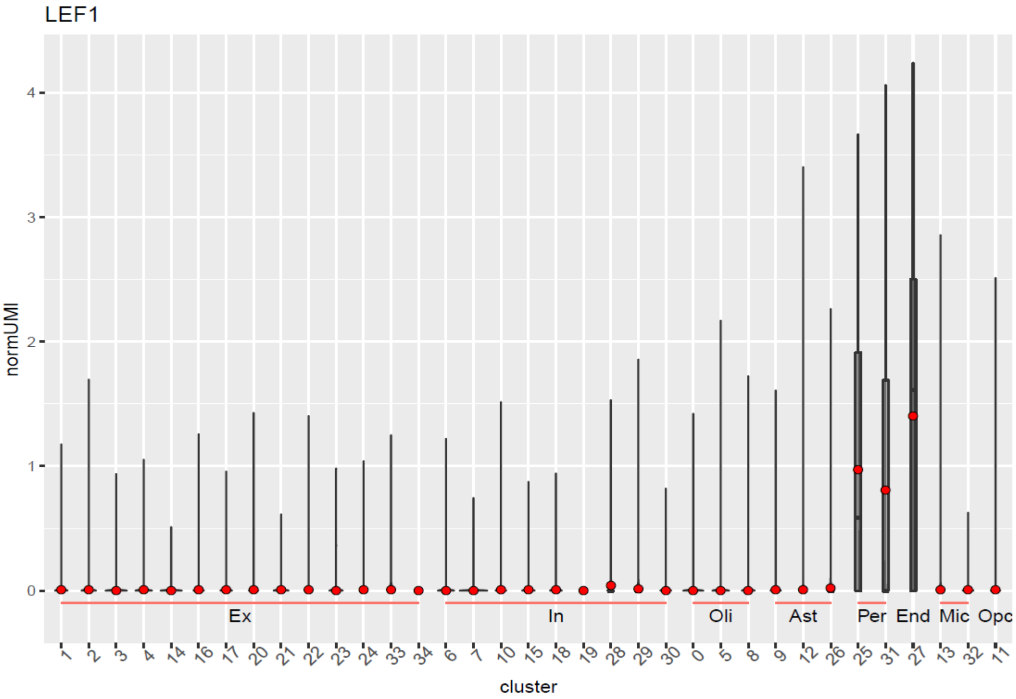


**Figure S7: Signature genes in brain vascular nuclei clusters. A.** Expression of signature genes in vascular clusters. Cluster 25 (cl.25)=activated pericyte, cl.27=endothelia, cl.31=resting pericyte clusters. **B.** Only vascular nuclei clusters demonstrated expression of *LEF1.*

**Figure S8:**

| Endothelial cluster | Top 5 Enriched GO terms (q <0.05) of Signature Genes |
|---|---|
| cl.31 | EXTRACELLULAR_MATRIX_STRUCTURAL_CONSTITUENT<br>EXTRACELLULAR_MATRIX_COMPONENT<br>EXTRACELLULAR_MATRIX<br>EXTRACELLULAR_STRUCTURE_ORGANIZATION<br>CIRCULATORY_SYSTEM_DEVELOPMENT |
| cl.25 | G_PROTEIN_COUPLED_RECEPTOR_SIGNALING_PATHWAY<br>GLOMERULUS_DEVELOPMENT<br>RHO_PROTEIN_SIGNAL_TRANSDUCTION<br>COLLAGEN_TYPE_IV_TRIMER<br>COMPLEX_OF_COLLAGEN_TRIMERS |
| cl.27 | GROWTH_FACTOR_BINDING<br>ENDOTHELIUM_DEVELOPMENT<br>ENDOTHELIAL_CELL_DEVELOPMENT<br>CARDIOVASCULAR_SYSTEM_DEVELOPMENT<br>ANATOMICAL_STRUCTURE_FORMATION_INVOLVED_IN_MORPHOGENESIS |

**Figure S8: Top Enriched GO terms of signature genes in each vascular cluster.** Cl.31=resting pericytes, cl.25=activated pericytes, cl.27=endothelia.

**Figure S9:**

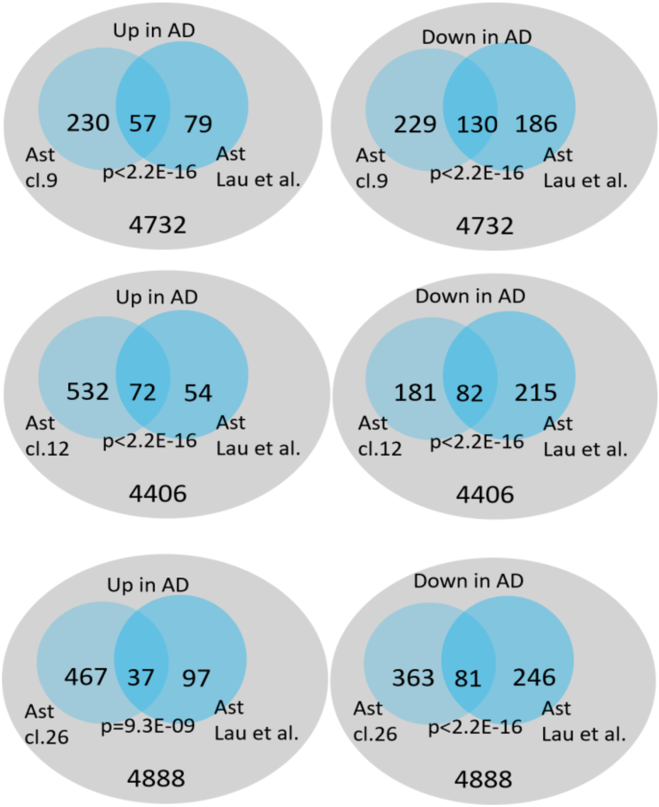


**Figure S9:** Number of overlapping DEGs between astrocytic clusters in our study (cl.9, cl.12 and cl.26) and those from Lau et al. study[1].

**Figure S10:**

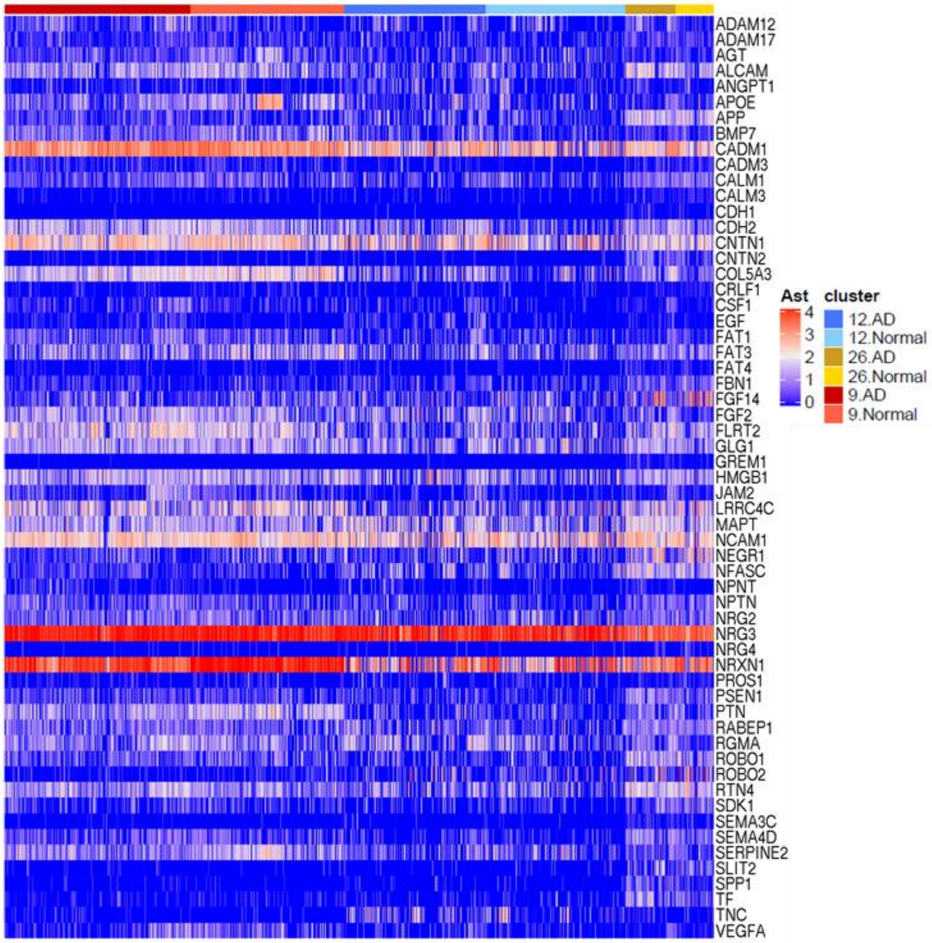


**Figure S10:** Heatmap of 59 astrocyte ligand genes identified through ligand-target interaction analysis platform Nichenet[2]. This analysis used significant DEGs between AD and control tissue identified in astrocyte clusters (cl.9, cl.12 and cl.26) as ligands and those in endothelial clusters (cl.25, cl.27 and cl.31) as potential targets. For more detailed description of this analysis, please see main text and method. Columns are cells arranged by clusters and diagnosis. Values shown in the heatmap are normalized expression.

**Figure S11:**

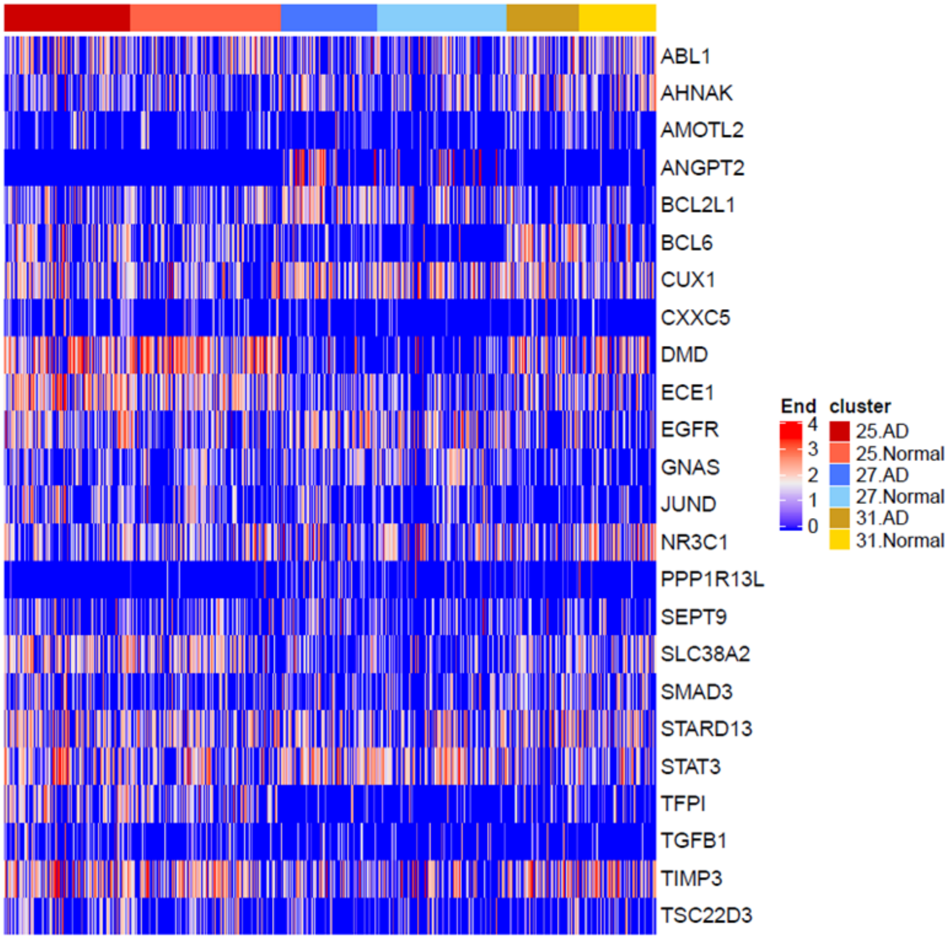


**Figure S11:** Heatmap of 24 vascular target genes identified through ligand-target interaction analysis platform Nichenet[2]. This analysis used DEGs between AD and control tissue identified in astrocyte clusters (cl.9, cl.12 and cl.26) as ligands and those in endothelial clusters (cl.25, cl.27 and cl.31) as potential targets. For more detailed description of this analysis, please see main text and method. Columns are cells arranged by clusters and diagnosis. Values shown in the heatmap are normalized expression.
